# Putative ligand binding sites of two functionally characterized bark beetle odorant receptors

**DOI:** 10.1101/2020.03.07.980797

**Authors:** Jothi K. Yuvaraj, Rebecca E. Roberts, Yonathan Sonntag, Xiaoqing Hou, Ewald Grosse-Wilde, Aleš Machara, Bill S. Hansson, Urban Johanson, Christer Löfstedt, Martin N. Andersson

## Abstract

Bark beetle behavior is to a large extent mediated via olfaction. Targeting the odorant receptors (ORs) may thus provide avenues towards improved pest control during outbreaks. Such an approach requires information on the function of receptors and their interactions with ligands. Hence, we annotated 73 ORs from an antennal transcriptome of the spruce bark beetle *Ips typographus* and report the functional characterization of two ORs (ItypOR46 and ItypOR49), which are selective for single enantiomers of the common bark beetle pheromone compounds ipsenol and ipsdienol, respectively. We use homology modeling and molecular docking to predict their binding sites. The importance of residues Tyr84 and Thr205 in ItypOR46 in the activation by ipsenol is experimentally supported, and hydrogen bonding appears key in pheromone binding. The biological significance of the characterized ORs positions them as prime targets for pest control and use in biosensors to detect bark beetle infestations.

## Introduction

Conifer-feeding bark beetles (Coleoptera; Curculionidae; Scolytinae) pose serious threats to forestry, and bark beetle outbreaks are increasing due to climate change^1–3^. Abiotic factors are important drivers for outbreaks^3^, with a warmer and drier climate reducing the defenses of the trees, at the same time as the beetles’ populations increase due to decreased generation time, lower winter mortality, and higher availability of breeding material resulting from severe weather events^2^. In light of the intensifying outbreaks, more efficient control and detection of bark beetles are needed. One avenue forward is to exploit their odorant receptors (ORs)^4^, which detect olfactory information crucial for successful mate and host finding and used to coordinate attacks on trees.

Neuronal responses that ultimately may induce a behavior are triggered in the olfactory system when odorants interact with ORs, which are located in the dendrites of olfactory sensory neurons (OSNs) in the antennae^5^. Insect ORs, which are unrelated to G-protein coupled vertebrate ORs^6,7^, are encoded by a large gene family^8^, undergoing a dynamic ‘birth-and-death’ evolution. In this model, gene duplication represents the birth, and pseudogenization and deletion the death of genes^9,10^. With few exceptions (reviewed in ^9^), a single OR gene is expressed in each OSN together with the co-receptor Orco, which is conserved across insects, except in the most basal taxa^11^. Together, the OR and Orco are suggested to form a heterotetrameric receptor complex^12^, with Orco being essential for the formation of an ion channel upon ligand-induced activation of the OR^13,14^. Current knowledge of the molecular and functional evolution of the ORs as well as the ligand-OR interaction is however limited, yet crucial for understanding insect chemical ecology and species-specific sensory adaptations. From an applied perspective, functional characterization of ORs and determination of their binding sites are pertinent in pest insects, because receptors that are key to survival and reproduction represent potential targets for improved pest control using OR antagonists and agonists^4^. Also, with the advancement of biosensor technology towards using ORs to detect insect semiochemicals^15–17^, employing receptors tuned to the characteristic odors of a pest, such as bark beetle aggregation pheromones, could be useful for early detection of infestations.

The European spruce bark beetle (*Ips typographus* L.) is the most serious pest of Norway spruce (*Picea abies* (L.) H. Karst.) in large parts of Europe and Asia^18^. As other bark beetles, *I. typographus* is a keystone species in forest ecosystems, contributing to the decomposition of wood through direct feeding as well as through the spread of microorganisms, such as its associated fungi^19^. When beetle populations reach a critical threshold, healthy trees are killed through mass-attacks, and entire forest landscapes can be quickly transformed. Attacks on trees are coordinated via an aggregation pheromone (*cis*-verbenol and 2-methyl-3-buten-2-ol) and attracts both sexes to trees^20^. Other compounds are released by the beetles during the later attack phases^21^, including verbenone, ipsenol, and ipsdienol, and several of them reduce attraction to the aggregation pheromone^22^. Most of these compounds are also used by other bark beetle species, frequently as aggregation pheromones or pheromone antagonists. Moreover, volatiles released from host trees^23^, non-host plants^24^, heterospecific bark beetles^25^, and fungal symbionts^19^ affect the behavior of *I. typographus*. Significant efforts to characterize OSN responses of *I. typographus* have been undertaken, with 23 strongly responding OSN classes reported, including neurons tuned to bark beetle pheromones, host and non-host volatiles, or fungal compounds^19,26–31^. The comparatively large knowledge of physiologically active compounds makes this species a good model for pursuing functional characterization of ORs.

Functional characterization of insect ORs has been biased towards moths^32–34^, flies and mosquitos^35,36^. In contrast, Coleoptera – arguably the largest order of the Metazoa– remains an understudied group, with only three characterized pheromone receptors in the cerambycid *Megacyllene caryae*^37^. For *I. typographus*, a previous study reported 43 ORs (ItypORs) from an antennal transcriptome^38^. The majority of these were, however, only reported as partial genes, and Orco was not identified. Here, we report a highly-improved complement of ORs in *I. typographus*, which allowed us to pursue functional characterization. Using two systems for heterologous expression and a panel of 68 compounds, we report the characterization of two ORs (ItypOR46 and ItypOR49), which are narrowly tuned to single enantiomers of the common bark beetle pheromones ipsenol and ipsdienol, respectively. To gain insight into the mechanisms of ligand binding in these ORs, we took advantage of the recently published cryo-EM structure of Orco^12^ to perform homology modeling and ligand docking simulations. This analysis predicted a primarily hydrophobic cavity lined by residues that are likely to interact with the ligands. The functional importance of two residues was supported experimentally using site-directed mutagenesis. The deorphanization of the two ItypORs and prediction of their ligand binding sites provide new insight into the interaction between insect ORs and their ligands, and represent important steps towards improved control of *I. typographus*, as this information may guide screenings for more potent agonists or antagonists. Future applications may also involve the use of these ORs in biosensors to detect a large number of bark beetle pests due to the widespread use of ipsenol and ipsdienol in bark beetle chemical ecology.

## Results

### OR annotation and phylogenetic analysis

To obtain an improved set of ItypOR sequences and to identify the Orco necessary for functional characterization in heterologous *in vitro* systems, we sequenced, assembled, and annotated an antennal transcriptome of *I. typographus*. The annotation revealed 73 ItypORs (including ItypOrco) of which 52 ORs corresponded to full-length proteins. Five of the original 43 ORs^38^ were discarded as previous assembly isoforms. Hence, 35 ORs were novel sequences. The majority of the previously partial OR sequences were extended to full-length, and sequences that contained e.g. previously unnoticed frameshifts or 5’/3’ intron sequence were corrected (Supplementary Table 1). The currently still partial ItypOR sequences encode proteins comprising 76 to 377 amino acids. Molecular cloning from cDNA followed by sequence verification allowed us to combine short partial OR sequences encoded by eight transcripts into four unique longer, but still partial, genes (ItypOR57NTE, ItypOR61NTE, ItypOR70FN, and ItypOR71NTE; gene suffixes are explained in the Methods section).

A recent study defined nine major clades of coleopteran ORs^39^. Our phylogenetic analysis of ORs from the Curculionidae and Cerambycidae families showed that the largest number of ItypORs were found in group 7 (29 ORs), followed by group 5A (21 ORs), group 1 (11 ORs), group 2A (7 ORs) and 2B (4 ORs), which is similar to the OR distribution in the other bark beetle species in the analysis, *Dendroctonus ponderosae* (Figure 1). The largest lineage expansions of ItypORs were present in groups 5A (ten ORs, but with low support) and 7 (seven ORs with 100% support). Additionally, *I. typographus* lacked ORs in groups 3, 4, 5B, and 6, which is also true for *D. ponderosae*. In contrast to previous studies^39,40^, our tree did not recapitulate the monophyly of OR group 2, with the two group 4 OR members from the cerambycid *Anoplophora glabripennis* being associated with the 2B group. This discrepancy is likely explained by the limited number of group 4 ORs in the analysis, in combination with the comparatively low node support for the 2A/2B distinction observed previously^39^. Our analysis also indicated the presence of 19 simple 1:1 orthologous relationships between ItypORs and DponORs of which 17 have high bootstrap support (≥90%; Figure 1). Apart from Orco, no simple orthologous relationships were observed between *I. typographus* and *A. glabripennis.*

**Figure 1.**
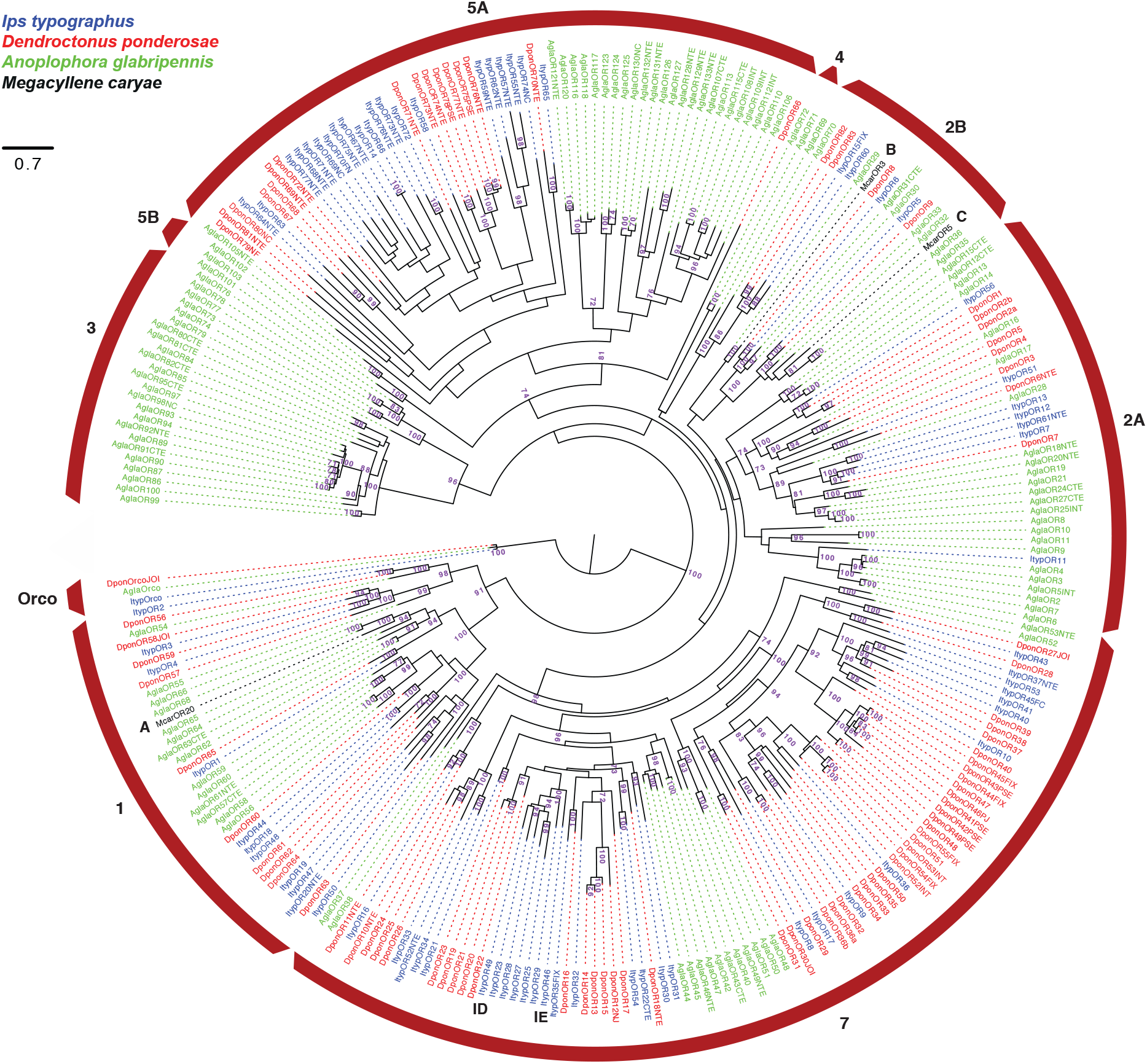
Maximum likelihood tree of beetle odorant receptors (ORs). The tree is based on a MAFFT alignment of amino acid sequences, constructed using RaXML, and rooted with the Orco lineage. Included are ORs from *Ips typographus* (Ityp; blue), *Dendroctonus ponderosae* (Dpon; red), *Anoplophora glabripennis* (Agla; green), and the three functionally characterized ORs from *Megacyllene caryae* (Mcar; black). Major coleopteran OR clades are indicated by the red arcs and numbered according to^39^. Numbers at nodes represent bootstrap support (n = 100), calculated using RaXML, and are only shown if ≥ 70. Key ligands for functionally characterized ORs indicated in the tree (data for McarORs from^37^): **A** = (2*S*,3*R*)-2,3-hexanediol (McarOR20); **B** = (*S*)-2-methylbutan-1-ol (McarOR3); **C** = 2-phenylethanol (McarOR5); **IE** = (*S*)-(−)-ipsenol (ItypOR46); **ID** = (*R*)-(−)-ipsdienol (ItypOR49). The sources of sequence data and explanation of receptor suffixes are detailed in the Methods section.

### Functional characterization of ItypOrco, ItypOR46, and ItypOR49

ItypOrco and ORs were transfected into inducible TREx/HEK293 cells for functional characterization. Cells stably expressing ItypOrco responded dose-dependently to the Orco agonist VUAA1 (Figure 2), representing the first example of VUAA1 responses in a beetle Orco protein. This cell line was transfected with each of ItypOR46 and ItypOR49. These two ORs share 43.4 % amino acid identity and are part of the same radiation within OR group 7, and hence evolutionary unrelated to the characterized pheromone receptors of *M. caryae* (Figure 1). The stably expressing ItypOrco/ItypOR46 and ItypOrco/ItypOR49 cell lines were analyzed by Western blot, which showed protein expression of myc-tagged ItypOrco and each of the two V5-tagged ItypORs. Proteins were detected in cells induced to express the exogenous Orco and OR genes, and not in the non-induced control cells, demonstrating proper regulation by the repression system (Supplementary Figure 1).

**Figure 2.**
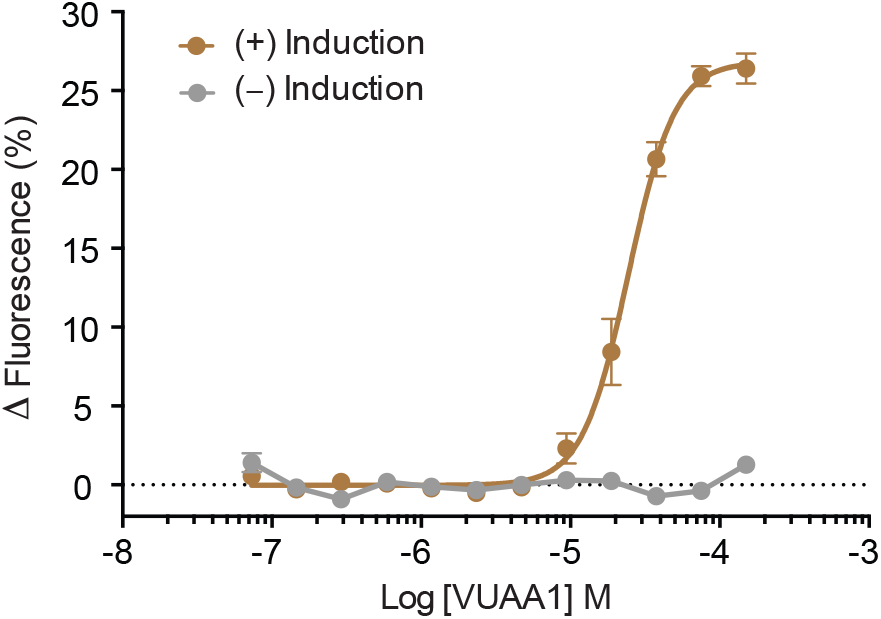
Dose-dependent response to the Orco agonist VUAA1 in TREx/HEK293 cells expressing ItypOrco. Data represent mean responses ± SEM (n = 3 biological replicates, each including 3 technical replicates, i.e., n_total_ = 9). EC_50_ of VUAA1 = 24.46 μM. (+)-Induction: response of cells induced to express ItypOrco; (−)-Induction: response of non-induced control cells.

Cells expressing ItypOrco/ItypOR46 and ItypOrco/ItypOR49 were screened for responses in a calcium fluorescence assay^41^ against a panel of 68 ecologically relevant compounds (Supplementary Table 2) at 30 μM concentration. In this experiment, ItypOR46 responded specifically to the pheromone compound (±)-ipsenol, with responses only recorded from induced cells (General Linear Model: F_1,14_ = 786; p < 0.001; Figure 3A, Supplementary Figure 2). A tendency for a secondary response to racemic ipsdienol was observed, but the response in the induced cells was not higher than that in non-induced cells (F_1,14_ = 1.17; p = 0.297). Dose-response trials with racemic ipsenol and its two pure enantiomers, which were synthesized in the present study (Supplementary Methods), showed that ItypOR46 is highly specific for the natural enantiomer (*S*)-(−)-ipsenol, with responses elicited by (*R*)-(+)-ipsenol only occurring at higher concentrations (Figure 3B). The response to (*R*)-(+)-ipsenol was likely due to the small percentage of the (*S*)-(−)-enantiomer present in the (*R*)-(+)-stimulus (Supplementary Table 2).

**Figure 3.**
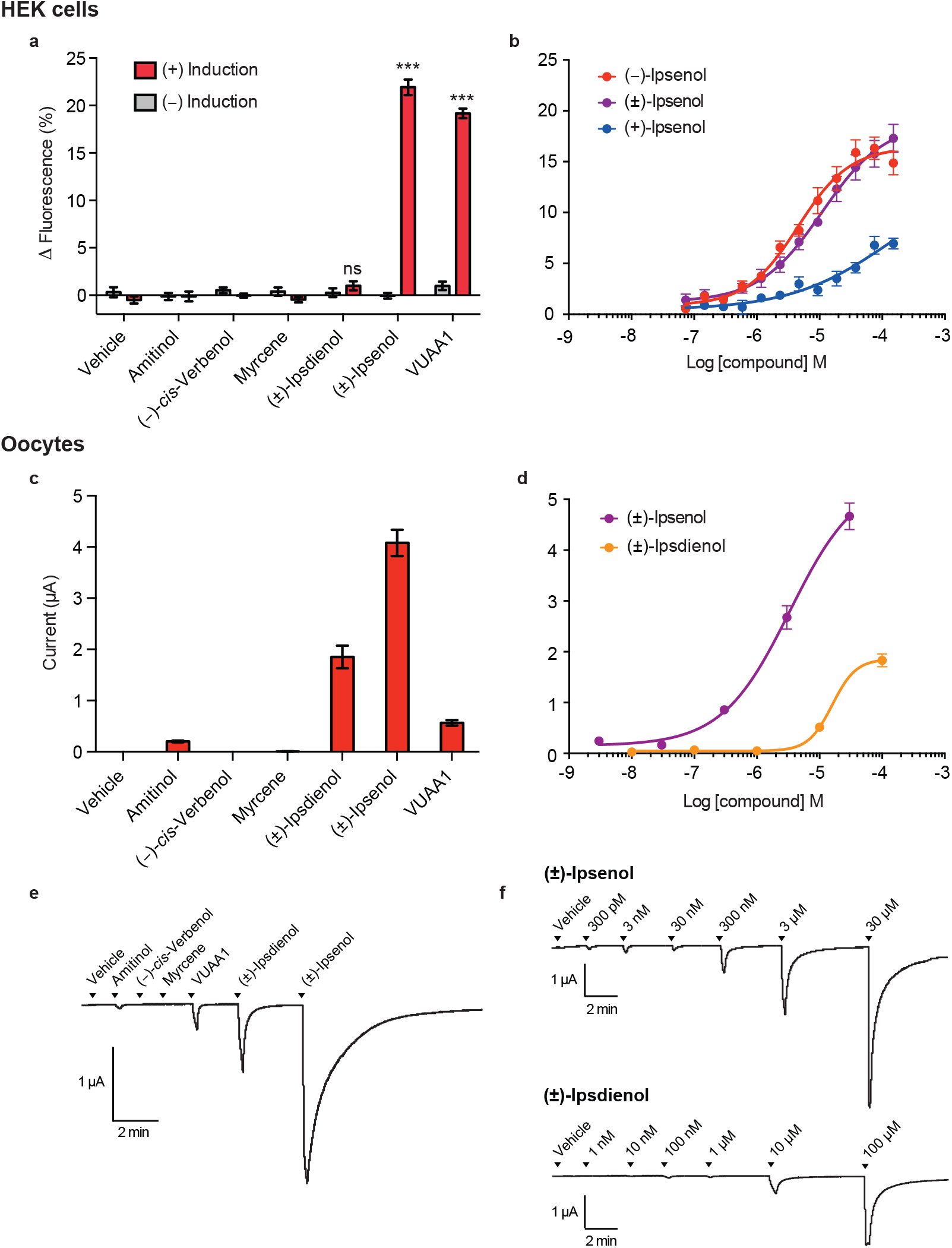
Functional characterization of ItypOR46. **a)** Response of TREx/HEK293 cells expressing ItypOR46 and ItypOrco to select stimuli (30 μM) and vehicle control in the screening experiment (responses to all 68 odor stimuli are shown in Supplementary Figure 2). (+)-Induction: response of cells induced to express ItypOrco and ItypOR46; (−)-Induction: response of non-induced control cells. Asterisks (***) indicate a significant difference (at p < 0.001) between induced and non-induced cells; ns = not significant (n = 3 biological replicates, each including 3 technical replicates, i.e., n_total_ = 9). **b)** Dose-dependent responses of the same TREx/HEK293 cell line to the pure enantiomers of ipsenol and the racemate (n = 4 biological replicates, each including 3 technical replicates, i.e., n_total_ = 12). EC_50_ values: (*S*)-(−)-ipsenol 1.98 μM; (±)-ipsenol 9.06 μM. **c)** Current responses of *Xenopus* oocytes expressing ItypOR46 and ItypOrco in the screening experiment to the same stimuli (30 μM) as shown in A) (n = 5). **d)** Dose-dependent current response of oocytes expressing ItypOR46 and ItypOrco to racemic ipsenol (n = 6) and ipsdienol (n = 5). **e)** Example of current trace responses from an oocyte expressing ItypOR46 and ItypOrco in the screening experiment (30 μM stimulus concentration). **f)** Example of current trace responses from two oocytes expressing ItypOR46 and ItypOrco in dose-response experiments. Upper and lower traces show responses to racemic ipsenol and ipsdienol, respectively. Data represent mean responses ± SEM (panels a-d).

Because insect OR responses sometimes depend on the system used for functional characterization^42^, the ORs were also tested in *Xenopus* oocytes using a reduced odor panel of six compounds (30 μM). Ipsenol was still the most active ligand, but ipsdienol elicited a secondary response in this system (Figure 3C, E). A minor response was elicited also by the structurally similar compound amitinol, but this response could well be attributed to the presence of ipsdienol (3%) in the stimulus, an impurity which was identified by GC-MS. The dose-response trials in the oocyte system indicated a higher sensitivity of ItypOR46 towards racemic ipsenol compared to ipsdienol (Figure 3D, F).

HEK cells expressing ItypOrco/ItypOR49 responded specifically to racemic ipsdienol in the screening experiment (F_1,14_ = 28.02; p < 0.001; Figure 4A, Supplementary Figure 3). A tendency for a secondary response to racemic ipsenol was observed, but it was not higher in induced compared to non-induced cells (F_1,14_ = 1.61; p = 0.225). Dose-response experiments that included racemic ipsdienol and its two pure enantiomers (synthesized in the present study; Supplementary Methods) showed that ItypOR49 is specifically tuned to (*R*)-(−)-ipsdienol, with responses to the (*S*)-(+)-enantiomer occurring only at higher stimulus concentrations. As with ItypOR46, these responses were likely due to the low percentage of (*R*)-(−)-ipsdienol in the (*S*)-(+)-enantiomer stimulus (Supplementary Table 2). ItypOR49 was generally non-responsive in the oocytes, apart from minute ipsdienol-induced changes in current (approx. 5 nA) in occasional oocytes (Supplementary Figure 4).

**Figure 4.**
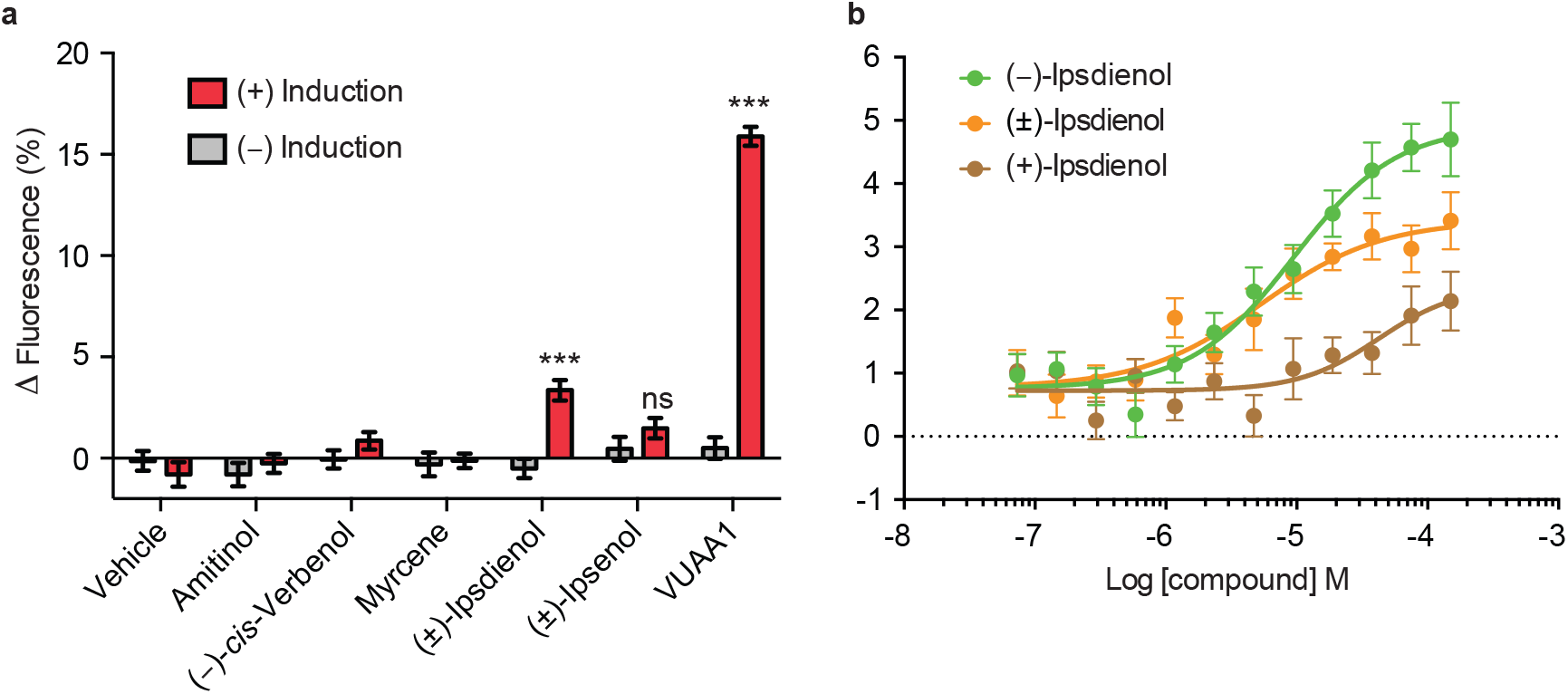
Functional characterization of ItypOR49. **a)** Response of TREx/HEK293 cells expressing ItypOR49 and ItypOrco to select stimuli (30 μM) and vehicle control in the screening experiment (responses to all 68 odor stimuli are shown in Supplementary Figure 3). (+)-Induction: response of cells induced to express ItypOrco and ItypOR49; (−)-Induction: response of non-induced control cells. Asterisks (***) indicate a significant difference (at p < 0.001) between induced and non-induced cells; ns = not significant (n = 3 biological replicates, each including 3 technical replicates, i.e., n _total_ = 9). **b)** Dose-dependent responses of the same TREx/HEK293 cell line to the pure enantiomers of ipsdienol and the racemate (n = 6 biological replicates, each including 3 technical replicates, i.e., n _total_ = 18). EC_50_ values: (*R*)-(−)-ipsdienol 9.47 μM; (±)-ipsdienol 5.34 μM. Data represent mean responses ± SEM.

As expected, significant responses to the Orco agonist VUAA1 were recorded from HEK cells expressing ItypOrco/ItypOR46 (F_1,14_ = 792; p < 0.001; Figure 3A) and ItypOrco/ItypOR49 (F_1,14_ = 469; p < 0.001; Figure 4A), indicating functional Orco expression. VUAA1 responses were also recorded from oocytes co-injected with ItypOrco and each of the two ORs (Figure 3C, E, Supplementary Figure 4). In HEK cells expressing ItypOrco/ItypOR46, the VUAA1 response magnitude at the 30 μM concentration was similar to the response elicited by racemic ipsenol (Figure 3A), whereas the VUAA1 response of Orco in the oocytes was 7-fold lower than the response to ipsenol (Figure 3C, E).

### Protein modeling and molecular docking

To gain insight into the ligand binding mechanisms of ItypOR46 and ItypOR49, protein homology modeling and molecular docking simulations were performed. First, an alignment of the two ItypORs with 3,185 additional ORs and Orco sequences^43,44^ was generated (Supplementary Data 1). Key residues with high conservation among ORs and Orco proteins were used to assess correct alignment and threading of the modeled ORs. The models of ItypOR46 and ItypOR49 revealed a putative binding cleft, exposed to the extracellular side. Several residues that have been implied to affect ligand specificity in various ORs (reviewed in ^44^) line this cleft (Figure 5), and significant differences between ItypOR46 and ItypOR49 were observed (described below), which may account for their dissimilarities in ligand specificity. Residues that when mutated were shown to affect inhibition of odor detection by DEET are located on transmembrane helix 2^ref45^, and residues affecting 2-heptanone specificity and pheromone activity^46,47^ are located on transmembrane helix 3, extra cellular loop 2 and transmembrane helix 4. The locations of these residues indicate that they possibly affect ligand binding (Figure 5), both at the extracellular entrance and at the deep end of the cleft, approximately at the center of the transmembrane domain. Hence, the docking site was defined to explore OR-ligand interaction possibilities throughout this confined region.

**Figure 5.**
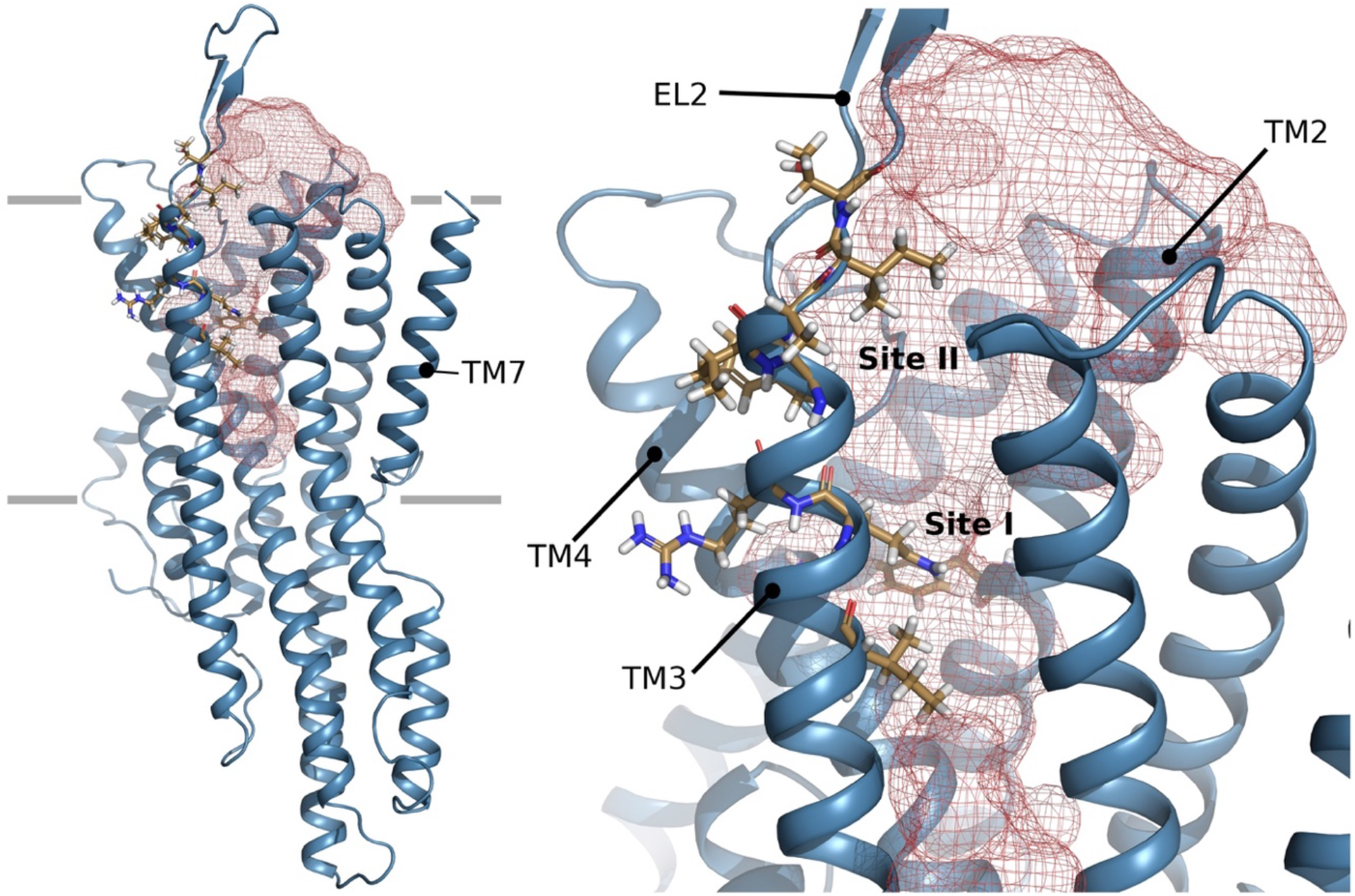
Protein model of ItypOR49. *Left image:* overview model indicating the predicted binding cavity in red mesh. Residues lining this cavity which in previous studies have been shown to affect ligand binding are showed in stick representation. Transmembrane helix 7 that forms the ion channel in the tetrameric Orco complex is located to the right. The expected extension of the lipid bilayer is indicated by grey lines. *Right image:* closer view of the cavity in the transmembrane region. Predicted binding site I and II as well as transmembrane (TM) domains 2-4 and extracellular loop (EL) 2 are indicated.

Molecular docking simulations included both enantiomers of ipsenol and ipsdienol, as well as the structurally similar but inactive compounds amitinol and myrcene, and the unrelated compound 1-hexanol (Supplementary Figure 5). Ipsenol and ipsdienol docked to two distinct locations in ItypOR46 and ItypOR49 respectively, directed by OR-specific residues that line the cleft. One deep site (site I) is located where mutational studies identified residues important for inhibition of odor detection by DEET^45^, and one site is closer to the extracellular opening of the cleft (site II; Figure 5). Most of the residues that have been shown to affect OR responses are concentrated at these two sites, or in their vicinity [reviewed in ^44^].

In ItypOR46, both enantiomers of ipsenol interacted via their conjugated double bonds in a π-π electron interaction with Tyr84, while being able to form a hydrogen bond to Thr205 as well as to Tyr84 at site I (Figure 6A). Hence, their poses were not sufficiently different to account for the enantiomer discrimination of the OR. The corresponding residues in ItypOR49 are Phe87 and Gly203. Therefore, both of these hydrogen bond interactions are absent in ItypOR49, and ipsenol was consequently instead found at site II, contacting helices 3 and 5 lined by Phe313 and hydrogen bonded to Tyr175 and Ser181 ((*R*)-(+)-ipsenol) or hydrogen bonded to Gln153 and lined by Phe313 and in a π-π interaction with Tyr175 ((*S*)-(−)-ipsenol) (Figure 6B).

**Figure 6.**
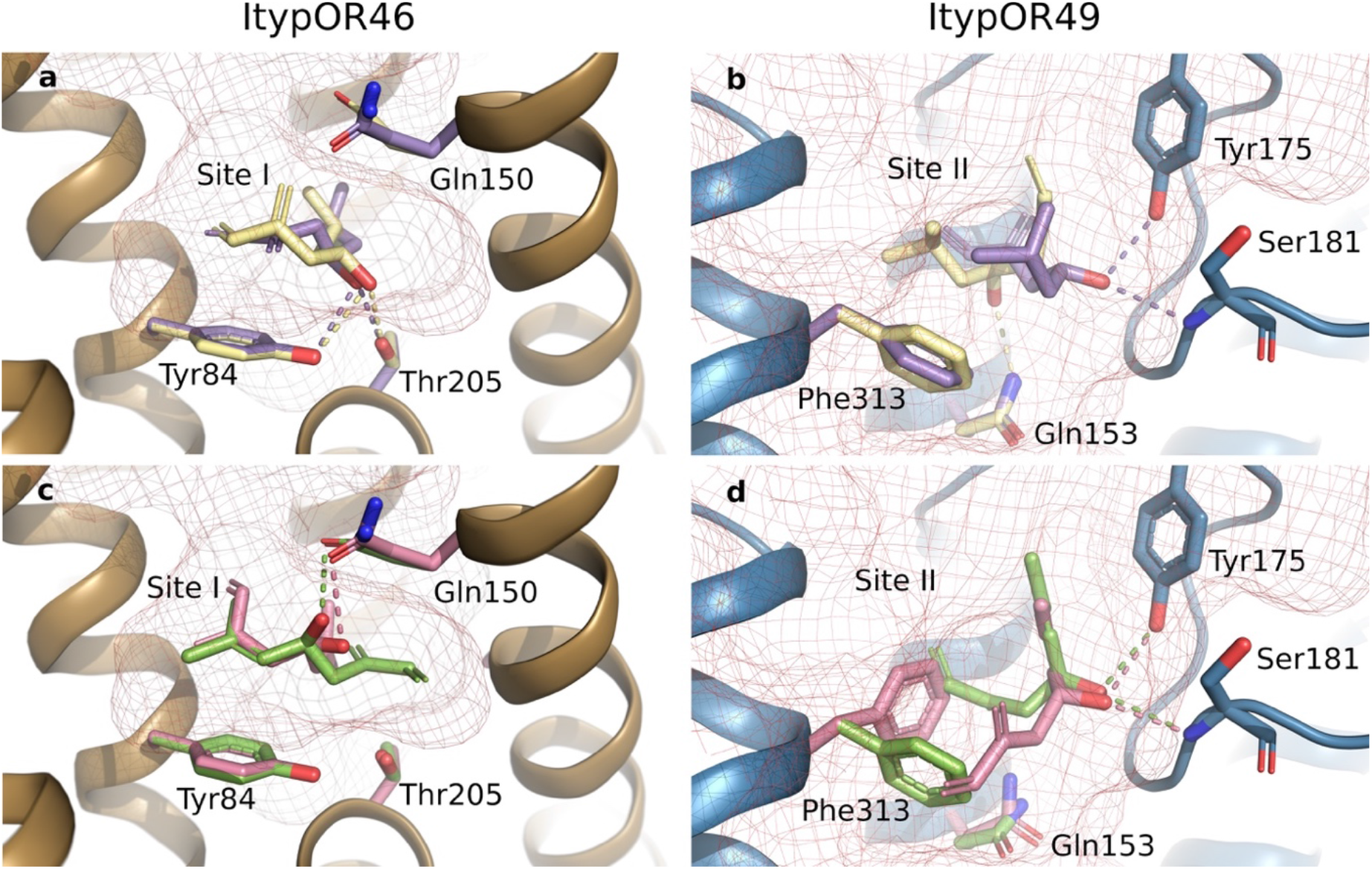
Results from molecular docking analysis. Predicted binding of (*R*)-(+)-ipsenol (purple) and (*S*)-(−)-ipsenol (yellow) to **a)** ItypOR46 (site I) and **b)** ItypOR49 (site II). Predicted binding of (*R*)-(−)-ipsdienol (rose) and (*S*)-(+)-ipsdienol (green) to **c)** ItypOR46 (site I) and **d)** ItypOR49 (site II). Binding site I is located approximately midway in the transmembrane region in relation to the plasma membrane. The shallower predicted binding site II is near the extracellular opening of the binding cavity. Potential hydrogen bonds are indicated with dashed lines. Residues in ORs that adopt a new position upon docking of a ligand are colored according to the corresponding ligand.

The same distribution between sites holds true for the ipsdienol enantiomers. In ItypOR46, the π-π interactions resulting from the docking simulations were similar to ipsenol π-π stacking to Tyr84, but no hydrogen bond to Thr205 at site I was observed. Instead, a hydrogen bond between the hydroxy group and Gln150 was favored (Figure 6C). Noteworthy, none of the 20 top poses of the ipsdienol enantiomers featured both a π-π stacking to the Tyr84 and a hydrogen bond to Thr205 in ItypOR46, substantially diminishing the favorable interaction as compared to the ipsenol enantiomers. Docking of the ipsdienol enantiomers into the ItypOR49 cleft resulted in binding to site II, involving π-π interactions with Phe313 for both enantiomers, hydrogen bond interactions to Tyr175 from extracellular loop 2 as well as to the backbone amine of Ser181 (Figure 6D). The favorable hydrogen bonds to the hydroxy group at the stereogenic center resulted in differing side chain rotamers and spatial occupancy of the ligand. As for ItypOR46, elucidating the enantiomer-specific activation of ItypOR49 requires knowledge of the conformation of the open ion pore state.

Poses of the inactive compound 1-hexanol did not cluster into any one specific site, but instead offered hydrogen bond donation to backbone carbonyls and hydrophilic side chains, while the extended hydrocarbon chain engaged in unspecific hydrophobic interactions. Myrcene engaged in unspecific hydrophobic interactions and in π-π electron stacking with either an aromatic moiety at the bottom of the cleft (Tyr84 in ItypOR46 and Phe87 in Ityp49) or with a pair of phenylalanines in helix 5 (ItypOR46: Phe316/Phe319; ItypOR49: Phe313/Phe317). Likewise, amitinol with its three double bonds, two of which are conjugated as in the myrcene structure, also interacted with the same aromatic residues in π-π stacking with the aforementioned residues. Having a tertiary alcohol group, amitinol is available for hydrogen bonding interactions, but in the confines of the binding cleft they are not equivalent to those available to the ipsenol and ipsdienol enantiomers.

### Site-directed mutagenesis of predicted ligand-binding residues

To gain support for the docking analysis, we introduced mutations to the two predicted key residues (Tyr84 and Thr205) at site I in ItypOR46, and used the HEK-cell assay to test the responses of mutated versions of the OR to the enantiomers of ipsenol. Because ipsenol was predicted to form a hydrogen bond with the hydroxy group present on the aromatic moiety of the tyrosine, we introduced two mutations to Tyr84: Tyr84Phe and Tyr84Ala. The former mutation removed the hydroxy group but retained the aromatic structure, whereas the latter mutation removed both features. Responses to (*S*)-(−)-ipsenol in cells expressing ItypOR46^Tyr84Phe^ were highly reduced as compared to responses in the wildtype receptor (included as control) and only observed at the highest concentrations (Figure 7A). In fact, the responses to (*S*)-(−)-ipsenol in this mutated OR were lower than the responses to (*R*)-(+)-ipsenol in the wildtype receptor. Cells expressing ItypOR46^Tyr84Ala^ did not respond to (*S*)-(−)-ipsenol at any concentration (Figure 7B), supporting the prediction of hydrogen bonding between the tyrosine and ipsenol at this site, but also suggesting that the presence of an aromatic residue has a minor importance. To investigate the functional importance of residue 205, we mutated this residue from Thr to Ala. Again, responses to (*S*)-(−)-ipsenol of cells expressing ItypOR46^Thr205Ala^ were completely abolished (Figure 7B). None of the mutated versions of ItypOR46 responded to (*R*)-(+)-ipsenol. Proteins of all mutated versions of ItypOR46 were detected by Western blot at equivalent or higher band intensities as the wildtype receptor, indicating sufficient levels of mutated proteins in the cells (Supplementary Figure 1).

**Figure 7.**
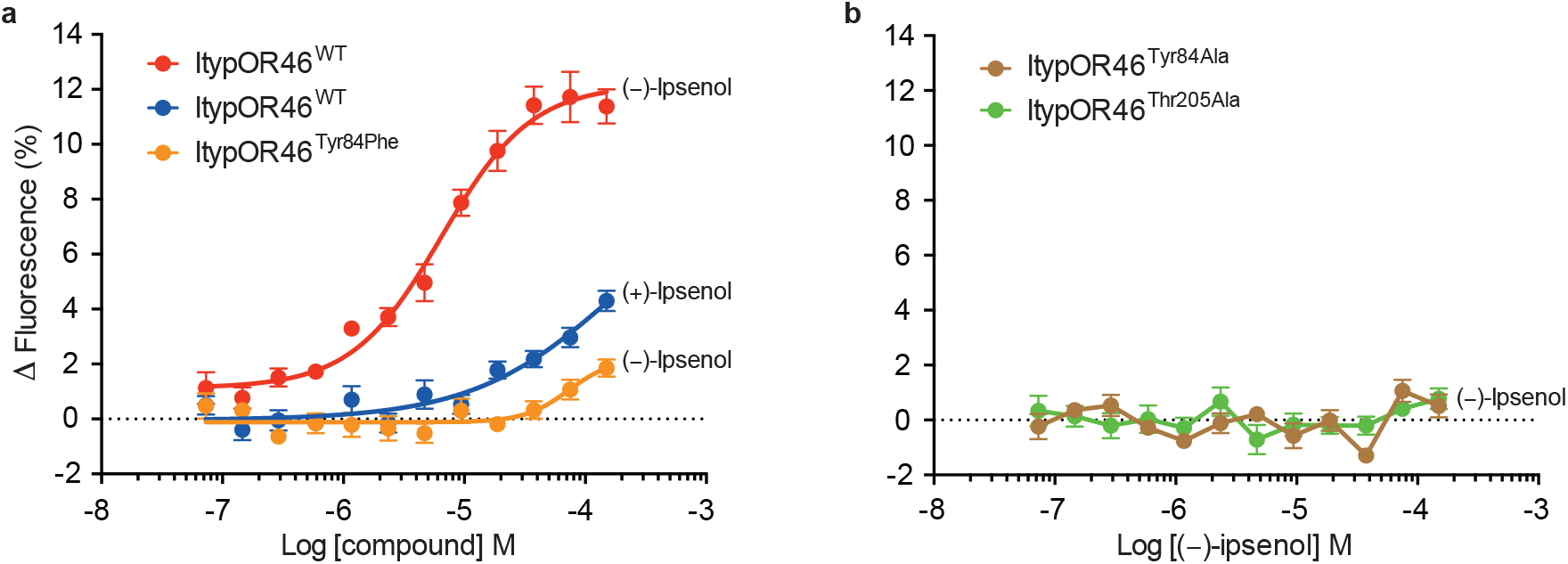
Response to ipsenol enantiomers of cells expressing wildtype (WT) ItypOR46 and mutated versions of this receptor. Data are split between two panels for clarity. **a)** ItypOR46^WT^ (positive control; n = 3 biological replicates, each including 3 technical replicates, i.e., n _total_ = 9) and ItypOR46^Tyr84Phe^ (n = 4 biological replicates, i.e., n _total_ = 12). **b)** ItypOR46^Tyr84Ala^ and ItypOR46^Thr205Ala^ (n = 3 biological replicates, n _total_ = 9 for both cell lines). For clarity, data for (*R*)-(+)-ipsenol are only shown for ItypOR46^WT^, since this compound did not activate mutated versions of the receptor. Data represent mean responses ± SEM.

## Discussion

We sequenced a new antennal transcriptome of *I. typographus*, and re-annotated its expressed ORs. The current dataset represents a marked improvement of the originally reported OR repertoire of this species^38^, including the identification of 35 novel ORs, 52 full-length ORs, and the essential co-receptor Orco. This, in turn, allowed us to perform functional characterization of the first bark beetle ORs and predict their binding sites, including amino acid residues with which the ligands are likely to interact. The identification of ItypOrco also allowed us to record the first responses of a coleopteran Orco to the agonist VUAA1^48^.

The number of ORs (73) in the *I. typographus* transcriptome is close to the number of putatively functional ORs (79) in the genome of the mountain pine beetle *D. ponderosae*^39^, and clearly higher compared to previous analyses of antennal transcriptomes of this and other species of bark beetles^38,49^, suggesting a good coverage of our analysis. The larger number of ORs compared to the original ItypOR dataset is likely due to improved sequencing depth and methodology^38^. A recent phylogenetic analysis of ORs from ten species across the Coleoptera identified nine major OR clades^39^. It showed that OR lineage radiations and losses among the nine clades differ remarkably between taxa, but also that OR distributions are more similar between closely related beetle families. Accordingly, both species of bark beetles included in our analysis have most of their receptors in OR group 7 and 5A, and both lack ORs from groups 3, 4, 5B, and 6. Similar distributions were reported also from other species of the Curculionidae family^49,50^. In contrast, the OR distribution in other beetle families, including the relatively closely related cerambycids (e.g., *A. glabripennis*)^51^, are different, with most species having representatives from the groups missing in bark beetles, and a lower proportion of ORs in group 7. Whether these differences are adaptive and relate to ecological specializations or if they represent chance evolutionary events^10^, remains unknown. The general lack of simple OR orthology across beetle families, however, suggests that convergent evolution is an important driver for the function of ORs, since unrelated beetle species often have OSNs specialized for the same compounds, such as green leaf volatiles^27,52^. The phylogenetic positions of the pheromone receptors ItypOR46 and ItypOR49 in relation to the positions of characterized receptors in *M. caryae* suggest that pheromone receptors in beetles do not cluster in specific clades as they do in Lepidoptera^34^.

The current ItypOR dataset forms an important platform for functional characterization. Among the comparatively large number of characterized OSN classes in this species^2,19,27–31^ are neurons that respond specifically to (*S*)-(−)-ipsenol and (*R*)-(−)-ipsdienol, respectively^28,30,31^. These enantiomers are produced by male *I. typographus* during the later attack phases, i.e., the production starts when males have admitted females, and production peaks when females have started to lay eggs^21^. Field experiments have shown a clear antagonistic effect of ipsenol on the attraction to the aggregation pheromone (“shut-off” signal for aggregation), whereas behavioral effects of ipsdienol are less clear and concentration dependent^22^. Here, we report the ORs that are responsible for the detection of these compounds. In HEK-cells, ItypOR46 responded exclusively to ipsenol, whereas a secondary response to ipsdienol was recorded in the oocytes. The OSNs tuned to (*S*)-(−)-ipsenol also showed a secondary response to ipsdienol, and our responses from the oocytes match the *in vivo* data^27,28^, whereas the HEK cell recordings suggest a higher specificity. The reason(s) for the system-dependent specificity of this receptor remains obscure, but may be due to differences between HEK cells and oocytes in their cell membrane compositions, affecting the folding of the receptor and access to the ligand binding site. System-dependent OR responses were recently documented for moth ORs^42^. The weak response to amitinol of ItypOR46 in oocytes is not reflected at the OSN level, and was likely due to the 3% ipsdienol impurity in the stimulus.

Our HEK-cell data for ItypOR49 showed a specific response to ipsdienol, but this response was weaker than the response of ItypOR46 to ipsenol. In contrast, the VUAA1 responses of Orco in these two cell lines were comparable. Our Western blot analysis revealed only a faint band for ItypOR49 (from two independent cell lines), which suggests that protein expression of this OR was low. The OSNs tuned to (*R*)-(−)-ipsdienol displayed a weaker response to ipsenol^27,30^, but this response was not observed for ItypOR49, which may be due to the overall low responses of this OR in the HEK cells. Similar to previous OSN recordings, our data reveal a high selectivity for enantiomers of ipsenol and ipsdienol in the two ORs, with apparent responses to the non-selected enantiomer being mostly explained by small enantiomeric impurities. Ipsenol and ipsdienol are produced, and have been shown to affect behavior, and/or trigger selective OSN responses in several *Ips* and *Dendroctonus* species^26,31,53^. Yet, orthologues of ItypOR46 and ItypOR49 have not been identified in the *Dendroctonus* species investigated so far^39,49^, again suggesting that convergent evolution is an important driver for OR function in beetles, even within a taxonomic subfamily. Whether the enantiomers of the two pheromone compounds are detected by orthologous ORs in other *Ips* species remains to be investigated by identifying ORs from additional species in this genus.

The recent structure of an Orco tetramer^12^ can be used in homology modeling of insect ORs, assuming that OR and Orco proteins fold similarly to adopt similar structures. This assumption is reasonable because ORs and Orco are believed to share a common ancestor^8^ and because structural features of proteins generally are more conserved than their functions. Similar to the Orco structure^12^, our models of ItypOR46 and ItypOR49 revealed a cleft exposed to the extracellular side. Based on its location, it is reasonable to assume that this cleft is important for ligand binding, and this assumption is further supported by numerous studies of other ORs where mutations to residues lining this cleft have affected the responses (summarized in ^44^). Our docking analysis towards this cleft suggested two discrete binding sites in ItypOR46 and ItypOR49, respectively. Apart from (*R*)-(+)-ipsenol and (*S*)-(+)-ipsdienol, the inactive compounds were not predicted to interact sufficiently with these residues. Ipsenol was predicted to interact with Tyr84 and Thr205 at site I in ItypOR46, whereas ipsdienol did not interact with the latter residue, which may relate to its lower activity on this OR. The corresponding residue to Tyr84 (Val91) in DmelOR59b has been shown to be central for the inhibitory effect of DEET on odor detection^45^. This residue is also adjacent to one of the sites that affects VUAA1 responses in Orco proteins from different species^43^. Interactions between enantiomers of the two active compounds to Gln150 in ItypOR46 and the corresponding Gln153 in ItypOR49 were also observed, and this residue has been shown to be important for the responses to 2-heptanone in DmelOR85b^54^. Based on its robust responses to ipsenol, we targeted ItypOR46 to provide experimental support for the predicted importance of Tyr84 and Thr205 in ligand binding. Mutating any of these residues to alanine completely abolished the response to ipsenol, whereas cells expressing ItypOR46^Tyr84Phe^ retained a small response at the highest stimulus concentrations. The latter response indicates that this mutant OR is likely correctly folded, although severely affected in the binding of ipsenol. Because the aromatic phenylalanine differs from tyrosine only by lacking the hydroxy group, the predicted hydrogen-bonding between ipsenol and Tyr84 was supported, suggesting that this interaction is crucial for the mechanism leading to opening of the ion channel of the receptor complex.

Opening of the ion channel upon binding of ligands is likely to involve a two-step mechanism – binding of the activating ligand, followed by a conformational change transmitted to helix 7, which is blocking the ion channel in the tetrameric Orco complex. The structure of the *Apocrypta bakeri* Orco^12^ revealed the closed structure, and as such is ideal for studying ligand binding sites. Theoretically, because opening of the ion channel results from ligand binding, it follows that the open state binding site must have higher affinity than the binding site at the closed state in order to drive the conformational change. Additionally, because the cleft is water filled and primarily hydrophobic, as revealed by the Orco structure and likewise our OR homology models, binding of aliphatic chain ligands could likely favor the open state by lowering the free energy barrier required for the transition as well, via expulsion of water molecules. Without a structure of the open channel, the underlying mechanism of channel opening upon binding of ipsenol and ipsdienol, and how the different enantiomers are discriminated by the ORs remain obscure. Our findings also raise questions regarding how ligand specificity may evolve in insect ORs. Ipsdienol differs from ipsenol only by the presence of an additional double bond. Yet, the two compounds are detected by ORs that only share 43.4% amino acid identity. Although all of this variation is unlikely to affect selectivity, this suggests that complex molecular changes may underlie specificity shifts in insect ORs detecting similar compounds. On the other hand, such major changes may be exactly what is required for ORs to display such a high discrimination for compounds being that similar. Further investigation is needed to understand the molecular evolution that determines ligand selectivity in insect ORs; in particular, revealing the structure of a ligand-binding OR would be especially rewarding.

Because ipsenol elicits strong antagonistic effects on pheromone attraction in *I. typographus*, ItypOR46 would be a prime target to employ in screenings aimed to identify more potent agonists than the natural ligand. Such a screening can now be directed towards the predicted binding site, and such agonists could to be used in the development of more efficient repellents for forest protection. Indeed, agonists that elicit ultra-prolonged activation of the carbon dioxide-sensitive neurons of mosquitos have been identified, with extended effects on host-seeking behavior^55^. Whether or not similar compounds can be identified for ItypOR46, and how effectively they will divert attacks, remains to be investigated. Receptor-targeted control using ORs that are specific for a pest are likely associated with less off-target effects as compared to compounds targeting the conserved Orco^4^. Additionally, the widespread production of ipsenol and ipsdienol across many species of bark beetles makes both ItypOR46 and ItypOR49 suitable candidates to be used in sensitive biosensors^16^ for detection of infestations of different bark beetles. Although there are technical challenges to overcome before such sensors can be used for airborne volatiles in a field situation, early detection of infestations and removal of attacked trees are crucial to limit bark beetle population growth, and hence outbreaks and economic loss.

## Methods

### Insect material and RNA isolation

*Ips typographus* individuals originated from a laboratory culture reared on Norway spruce (*P. abies*) logs, and were kindly provided by Prof. F. Schlyter. The antennae from 255 adults (males and females combined in an equal sex ratio) were homogenized using Tissue-tearor model 98370-365 (Bartlesville, OK, USA), and total RNA was isolated using the RNeasy Minikit (Qiagen, Hilden, Germany). This yielded 6.2 μg of high quality total RNA that was used for transcriptome sequencing and molecular cloning.

### Transcriptome sequencing, annotation, and phylogenetic analysis of ORs

DNase treated RNA was subjected to poly-A enrichment and library construction using a RNA-Seq v2 Library Preparation Kit (Illumina, San Diego, CA, USA), followed by 150 bp paired-end sequencing, performed on an Illumina HiSeq 3000 platform at the Max Planck-Genome-centre (Cologne, Germany). The sequencing yielded 31,622,325 paired-end reads post quality appraisal and initial read filtering using standard methods. Low-quality reads and adaptor sequences were removed. The high-quality reads were *de novo* assembled using the short reads assembly program Trinity version 2.3.2^ref56^ as well as a CLC Genomics Workbench version 10 (Qiagen, Carlsbad, CA, USA). The Trinity assembly yielded a total of 74,151 predicted ‘genes’ with their isoforms totaling 171,567 predicted ‘transcripts’ (i.e., on average 2.3 assembly isoforms per predicted gene) with average length of 1,487 bp and N_50_ = 3,373 bp. The CLC assembly resulted in 47,576 assembled contigs with an average length of 1,033 bp and N_50_ = 1,488 bp. The overall completeness of these assemblies was assessed using the Benchmarking Universal Single-Copy Orthologs (BUSCOv3.0.1; https://busco.ezlab.org/) tool performed against the Insecta odb9 dataset, which included 1658 reference genes^57^. This analysis indicated that the percentage of complete BUSCOs was 97.4 for the Trinity assembly and 80.1 for the CLC assembly, but also a higher percentage of duplicated BUSCOs in the Trinity assembly (Supplementary Table 3). The sequence reads have been deposited in the SRA database at NCBI under the BioProject accession number PRJNA602798.

Sequences of *I. typographus* ORs (ItypORs) were annotated through tBLASTn searches against the above-mentioned assemblies using query sequences from *I. typographus*, *D. ponderosae* (Curculionidae), *A. glabripennis* (Cerambycidae) and *Leptinotarsa decemlineata* (Chrysomelidae) ^38,58,59^. An *e*-value cut-off at 1.0 was used to account for the divergent nature of this gene family. All identified ItypORs were included in additional BLAST searches until all novel hits were exhausted. Due to the difference in assembly completeness, the majority of the ItypOR sequences were identified from the Trinity assembly, with only a few ORs being more complete in the CLC assembly (Supplementary Table 1). A few OR sequences could be extended or completed by joining overlapping sequences from the two current assemblies and/or the published assembly^38^. Short transcripts encoding partial OR sequences that did not overlap with other ItypOR sequences in multiple sequence alignments were discarded to ensure that all reported ORs were unique. The previously identified ORs (ItypOR1-43) retained their original names, and novel ORs were given names from ItypOR44 to ItypOR77 in the order they were identified. Some of the partial original OR sequences^38^ were here discarded as assembly isoforms, but their numbers were not recycled for any of the novel ORs to avoid confusion. Transcripts did not always encode full-length OR sequences. Hence, suffixes were added to gene names following established practice^40^, with NTE and CTE suffixes given to genes with the N-terminus or C-terminus missing, respectively. A FIX suffix was given to genes that were annotated on transcripts which were manually corrected using raw RNAseq reads or following the other current or previously published assemblies^38^. In cases where genes had multiple suffixes, one-letter abbreviations were used in combinations (i.e., N, C, and F).

The amino acid sequences of the ItypORs were aligned with the ORs from the genomes of *D. ponderosae* (Dpon)^40^ and *A. glabripennis* (Agla)^58^ using MAFFT 7.017^ref60^, implemented in Geneious software package 7.1.9. The three functionally characterized ORs from *M. caryae* (Mcar) (Cerambycidae)^37^ were also included to indicate their positions in the phylogeny. Pseudogenes and partial ORs below 200 amino acids from *A. glabripennis* were excluded to improve the alignment and reduce the size of the tree. Uninformative regions of the alignment were excised using trimAl v1.2^ref61^ with the following settings: similarity threshold 0, gap threshold 0.7, and minimum 25% conserved positions. Partition finder 2^ref62^ was used to select a model of evolution, with the best fit obtained for a JTT amino acid substitution matrix, a proportion of invariant sites, gamma distributed rate variation, and empirical equilibrium amino acid frequencies (JTT+I+G+F). These parameters were used to construct a maximum likelihood tree using RAxML 8.1.2^ref63,64^, with branch support calculated by rapid bootstrapping (N = 100). The tree was visualized, rooted with the Orco lineage, and color coded in FigTree 1.4.3. Final graphical editing was performed using Adobe Illustrator.

### First-strand cDNA synthesis and confirmation of OR sequences

First-strand cDNA was synthesized from 1 μg of DNase-treated antennal RNA using the ThermoScript RT-PCR system for First-Strand cDNA Synthesis (Thermo Fisher Scientific, Carlsbad, CA, USA), according to the manufacturer’s instructions, except for using both random hexamers and oligo-dT primers in the reaction. Multiple sequence alignments of the initial set of ItypORs annotated here indicated that eight OR fragments likely belonged to non-overlapping parts of the same four genes (ItypOR57NTE, ItypOR61NTE, ItypOR70FN, and ItypOR71NTE). Hence, PCR amplification from cDNA followed by Sanger sequencing of the PCR products were performed to verify these joins and to add internal DNA sequence that were missing on the transcripts (25-45 bp). These partial genes were amplified using Pfu Phusion Flash high-fidelity master mix (Thermo Fisher Scientific), and with the forward primer designed for the most N-terminal transcript and reverse primer for the most C-terminal transcript. The PCR products were resolved on 1% TAE agarose gels, and bands of expected length were excised and purified using the Wizard® SV Gel and PCR clean-up system (Promega). Sequencing PCR was performed using the purified PCR products, their gene-specific primers, and the BigDye® Terminator v1.1 Cycle Sequencing Kit (Thermo Fisher Scientific). Sanger sequencing was performed using an Applied Biosystems™ capillary 3500 Genetic Analyzer (Thermo Fisher Scientific) at the sequencing facility at the Department of Biology, Lund University.

### Molecular cloning of ItypOrco and ItypORs for functional characterization in HEK293 cells

Sequences of ItypOrco, ItypOR46, and ItypOR49 were amplified from antennal cDNA, using full-length gene-specific primers and the Pfu Phusion Flash high-fidelity master mix. The PCR products were purified as described above, and then included in a second PCR reaction, but this time using extended primers to add a 5’ NotI recognition site, a Kozak sequence (‘cacc’) and an N-terminal epitope tag (c-Myc for ItypOrco and V5 for ItypORs), as well as a 3’ ApaI recognition site. The PCR products were purified and then digested using NotI and ApaI restriction enzymes (NEB, Ipswich, MA, USA). The modified OR sequences were separated by gel electrophoresis, and bands of expected length excised, purified and ligated into the expression vectors pcDNA™4/TO (Orco) and pcDNA™5/TO (ORs) (all Thermo Fisher Scientific), followed by transformation into HB101 competent cells (Promega). Successful transformation was confirmed by colony PCR, and positive colonies were grown in LB broth overnight with ampicillin. Plasmids were extracted using the GeneJET Plasmid Miniprep kit (Thermo Fisher Scientific), and then Sanger sequenced. Plasmids with verified Orco or OR sequence were transformed into competent cells, positive colonies were identified by colony PCR, and then grown in LB broth overnight. Large quantities of purified plasmids were obtained using the PureLinkTM HiPure Plasmid Filter Midiprep kit (Thermo Fisher Scientific). The cloned sequences of ItypOrco, ItypOR46, and ItypOR49 have been deposited in GenBank under the accession numbers MN987209-MN987211.

ItypOR46 was subjected to site-directed mutagenesis to verify the functional importance of two residues (Tyr84 and Thr205) predicted (see below) to be central for ligand binding. Three point mutations (Tyr84Phe, Tyr84Ala, and Thr205Ala) were introduced individually to ItypOR46 in pcDNA5™/TO using the Q5 Site-Directed Mutagenesis kit (New England Biolabs) following the manufacturer’s instructions. Successful mutants were identified using Sanger sequencing, and large quantities of plasmids obtained as described above.

### Generation of inducible cell lines expressing ItypOrco and ItypORs, and confirmation of protein expression

HEK293 cells stably expressing ItypOrco and ItypORs were produced and cultured according to previously described methods^41^. Briefly, an isogenic, tetracycline repressor-expressing (TREx) cell line^41^ was transfected with pcDNA™4/TO/ItypOrco, and cultured using zeocin and blasticidin selection antibiotics (NEB). Afterwards, this TREx/ItypOrco-expressing cell line was tested in a fluorescent calcium assay (described below) against the Orco agonist VUAA1 (>98 % purity, Sigma-Aldrich), which directly activates the Orco protein in nearly all insect species tested to date^43,48,65^, for confirmation of functional Orco expression. The TREx/ItypOrco cell line was then used in separate transfections with pcDNA™5/TO/ItypOR46, pcDNA™5/TO/ItypOR49, and mutated versions of ItypOR46, and cultured as described above, but with the addition of the pcDNA5™/TO-specific selection antibiotic hygromycin (Gold Biotech). The resulting TREx/ItypOrco/ItypOR cell lines were analyzed for protein expression of myc-tagged Orco and V5-tagged ORs by Western blot. Cells were cultured, induced to express exogenous Orco and ORs and pelleted as previously described^41^, with non-induced cells included as controls. The protein extraction and blotting also proceeded according to previously described methods^34^.

### Functional characterization in HEK293 cells

Cells expressing ItypOrco alone, and ItypOrco in combination with each of the ItypORs, were tested for responses to compounds in the previously described fluorescent calcium assay^34,41,65^. Briefly, cells were plated into poly-D-lysine coated 96-well plates, and induced to express ItypOrco and ItypORs. Half of the wells were left non-induced to serve as a negative control. Prior to the assay, the wells were loaded with a calcium-sensitive fluorophore (Fluo4-AM, Thermo Fisher Scientific), after which cells were investigated for ligand-induced receptor activation using a FLUOstar Omega plate reader (BMG Labtech, Ortenberg, Germany). Cells were tested in triplicates (technical replicates) on each plate (biological replicate), and at least three independent replicates were conducted for each cell line and experiment. The cells in each well were subjected to a single stimulation, and then discarded.

Test compounds were diluted in DMSO and assay buffer as previously described ^41^, with the final DMSO concentration in the wells being 0.5%. The Orco agonist VUAA1 was included in the assays as a control (30 μM concentration) for functional Orco expression, and also in dose-response experiments with the ItypOrco cell line. Assay buffer with 0.5% DMSO in was included as a negative control (vehicle) in all assays. The two wildtype ItypOrco/ItypOR expressing cell lines were screened against a panel of 68 ecologically relevant compounds (30 μM concentration), including the pheromones from a variety of bark beetle species, host and non-host compounds, as well as compounds from the fungal symbionts of *I. typographus* (Supplementary Table 2). This odor panel comprised all known key ligands for the previously characterized OSN classes of this species and several of their secondary ligands^2,19,26–28^, along with a few bark beetle pheromone compounds with unknown activity in *I. typographus*. Compound purities were analyzed by gas chromatography-mass spectrometry (GC-MS) (Supplementary Table 2). Mean ligand-induced responses (± SEM) in ORs were calculated and graphed in GraphPad Prism 6 (GraphPad Software Inc., La Jolla, CA, USA). Ligands that elicited significantly stronger responses in induced versus non-induced cells were regarded as active. Hence, a General Linear Model analysis was performed using IBM SPSS statistics v.23 to identify active compounds, with ‘induction (yes/no)’ included as a fixed factor, and ‘plate number’ as a random factor to account for the variation between plates. Screening responses below 1% increased fluorescence were regarded as ‘no response’ because they were within the range of random variation of the assay. Active compounds eliciting responses above 3% increased fluorescence at the 30 μM screening concentration were included in subsequent dose-response experiments, which were also designed to elucidate the enantiomer-specificities of ItypOR46 and ItypOR49 (the synthesis of ipsenol and ipsdienol enantiomers is described in the Supplementary Methods). The three mutated versions of ItypOR46 were tested against the enantiomers of ipsenol. Half-maximal effective concentrations (EC_50_) with 95% confidence intervals were estimated using the non-linear curve fit regression function in GraphPad Prism.

### Functional characterization in Xenopus oocytes

ItypOR46 and ItypOR49 were also assayed using *Xenopus laevis* oocytes. Gene-specific primers for the two ORs and Orco, designed to include a flanking 5’ Kozak sequence (‘gccacc’) and 5’ and 3’ recognition sites (BamHI and XbaI for ORs; EcoRI and XbaI for Orco), were employed in PCR reactions using the OR-containing HEK cell expression vectors as templates. The PCR products were purified, digested and cloned into the pCS2+ expression vector as described above. Large quantities of plasmids containing verified inserts were obtained using the plasmid Maxi kit from Qiagen. The plasmids were linearized using NotI (Promega), and the linearized DNA was purified and transcribed into complementary RNA (cRNA) using the SP6 mMESSAGE mMACHINE® kit (Invitrogen, Carlsbad, CA, USA).

Oocytes were surgically removed from *X. laevis* frogs (purchased from University of Portsmouth, UK), and treated with 1.5 mg/ml collagenase (Sigma-Aldrich, St. Louis, MO, USA) in oocyte Ringer 2 solution (containing 82.5 mM NaCl, 2 mM KCl, 1 mM MgCl_2_, 5 mM HEPES, pH 7.5) at 20 °C for 15-18 minutes. Stage V-VII oocytes were co-injected with cRNAs from the ItypOrco and ORs (50 ng of each), and then incubated in Ringer’s buffer (96 mM NaCl, 2 mM KCl, 5 mM MgCl_2_, 0.8 mM CaCl_2_, 5 mM HEPES, pH 7.6) containing 550 mg/L sodium pyruvate and 100 mg/L gentamicin at 18 °C for at least 3 days. The previously described two-electrode voltage clamp electrophysiological set-up was used to record whole-cell inward currents from oocytes in good condition (3-6 days post-injection) at a holding potential of −80 mV^66^. Test compounds were applied to the oocyte chamber by means of a computer-controlled perfusion system at a rate of 2 ml/min for 20 s with extensive washing with Ringer’s buffer at 4 ml/min between stimulations. Data were collected and analyzed using Cellworks software (npi electronic GmbH, Tamm, Germany).

Due to the limited number of channels in the perfusion system, only six compounds were included: VUAA1, the HEK cell-active ligands ipsenol and ipsdienol and the structurally related compounds amitinol and myrcene, as well as the aggregation pheromone component (−)-*cis*-verbenol. The compounds were dissolved in DMSO and Ringer’s buffer to desired test concentrations and a final DMSO concentration of 0.1 %. Ringer’s buffer with 0.1 % DMSO served as a negative control. Compounds were screened for receptor activity at a concentration of 30 μM, and the active compounds were subsequently included in dose-response trials using additional oocytes.

### Protein modeling and ligand docking simulations

Sequence alignment of ItypOR46 and OR49 was performed against a multiple sequence alignment containing 3,185 OR and Orco sequences^43,44^ (Supplementary Data 1), including the sequence for the *A. bakeri* Orco for which a homotetrameric cryo-EM structure was recently published^12^. Homology models of the two ORs were produced in Swiss-Model^67^ with the *A. bakeri* Orco structure (PDB ID 6c70) as template. Extra cellular loop 2 which is absent in the Orco structure was built using the program SuperLooper2^ref68^. The resulting model structures were energy minimized using NAMD^69^.

Three-dimensional structures of the two enantiomers of ipsenol and ipsdienol, as well as myrcene, amitinol and 1-hexanol were produced, and AutoDockTools 1.5.6 (ADT) was used to convert the ligand structure files to AutoDock ligand format (pdbqt). The ItypOR46 and ItypOR49 homology models were likewise converted to pdbqt format. As in the Orco structure^12^, an approximately 20 Å deep wedge-shaped cavity at the extra cellular side, formed by helices 2 to 6, was identified. Residues lining the cavity were defined as flexible and input pdbqt files for AutoDock Vina 1.1.2^ref70^ were produced using ADT as well as a grid box for molecular docking simulation, covering the entirety of the putative binding cavity. AutoDock Vina 1.1.2 was used to perform molecular docking simulation, and the top 20 poses of each ligand were outputted.

## Acknowledgements

We thank Pavlina Kyjakova for measurement of chiral GC-MS spectra related to the synthesis of ipsenol and ipsdienol. We thank Fredrik Schlyter for providing biological material, and Dineshkumar Kandasamy, Rikard Unelius, Blanka Kalinová, Fredrik Schlyter, and Anna Jirošová for providing test compounds. SNIC is acknowledged for allocated computing time at LUNARC. We thank Richard Newcomb for scientific input on a previous version of the manuscript. This study was funded by the Swedish Research Councils FORMAS (grants #217-2014-689 and #2018-01444 to MNA) and VR (#2017-03804 to CL), the Crafoord foundation (MNA), the Carl Trygger foundation (MNA), the Royal Physiographic Society in Lund (RER and MNA), Stiftelsen Olle Engkvist Byggmästare (YS and UJ), and the Max Planck Society (BSH and EG-W).

## Author contributions

MNA conceived and coordinated the study. RER, JKY, XH, YS, EG-W, UJ, CL, and MNA contributed to experimental design. EG-W, BSH, and MNA coordinated the sequencing. EG-W performed transcriptome assembly and BUSCO analysis. MNA performed annotation and phylogenetic analysis of ORs. RER, JKY, XH, and MNA performed molecular work. RER and JKY performed HEK-cell culturing and experiments with contributions from MNA. XH performed oocyte recordings. YS and UJ performed protein homology modeling and docking analysis. AM synthesized enantiomers of ipsenol and ipsdienol. MNA analyzed the data statistically and drafted the manuscript with contributions from YS. All authors provided editorial and scientific input on the final version of the manuscript.

## Supplementary Material

**Supplementary Table 1.** Annotation details of the *Ips typographus* odorant receptors (ORs), including names, amino acid sequences, annotation notes, and correspondence with the original dataset^38^.

**Supplementary Table 2.** Compounds used for characterization of *Ips typographus* odorant receptors (ORs) and Orco, including their purities, source information, and examples of main biological origins.

**Supplementary Table 3.** Assessment of the completeness of antennal transcriptome Trinity and CLC assemblies using the Benchmarking Universal Single-Copy Orthologs (BUSCOv3) tool performed against the Insecta odb9 dataset (https://busco.ezlab.org/).

**Supplementary Figure 1.** Protein detection of *Ips typographus* odorant receptors (ORs; V5-tagged) and Orco (myc-tagged) from TREx/HEK293 cells by Western blot. *Upper left panel:* Detection of wildtype ItypOR46 and ItypOR49 (two cell lines). *Upper right panel:* detection of Orco in the same cell lines. *Lower panel:* detection of three versions of mutated ItypOR46 proteins and wildtype (WT) ItypOR46 (included as control). Proteins were only detected from cells induced (+) to express ItypORs and Orco, and not from non-induced (−) control cells, indicating proper regulation by the repression system.

**Supplementary Figure 2.** Response of TREx/HEK293 cells expressing ItypOR46 and ItypOrco to all stimuli (30 μM) and vehicle control in the screening experiment (n = 3 biological replicates, each including 3 technical replicates, i.e., n_total_ = 9). (+)-Induction: response of cells induced to express ItypOrco and ItypOR46; (−)-Induction: response of non-induced control cells. Data represent mean responses ± SEM.

**Supplementary Figure 3.** Response of TREx/HEK293 cells expressing ItypOR49 and ItypOrco to all stimuli (30 μM) and vehicle control in the screening experiment (n = 3 biological replicates, each including 3 technical replicates, i.e., n_total_ = 9). (+)-Induction: response of cells induced to express ItypOrco and ItypOR49; (−)-Induction: response of non-induced control cells. Data represent mean responses ± SEM

**Supplementary Figure 4.** Current traces of two oocytes expressing ItypOrco/ItypOR49, indicating responses to the Orco agonist VUAA1 and minute responses to racemic ipsdienol.

**Supplementary Figure 5.** Chemical structures and IUPAC names of compounds included in the molecular docking analyses against ItypOR46 and ItypOR49.

**Supplementary Methods.** Description of the chemical synthesis of ipsenol and ipsdienol as well as their pure enantiomers.

**Supplementary Data 1.** Multiple sequence alignment used to generate the ItypOR models.

