## Supplementary Figures, Tables and Methods for "Putative ligand binding sites of two functionally characterized bark beetle odorant receptors"

### Material included


| | |
|---|---|
| Supplementary Table 2 | S-2 |
| Supplementary Table 3 | S-3 |
| Supplementary Figure 1 | S-4 |
| Supplementary Figure 2 | S-5 |
| Supplementary Figure 3 | S-6 |
| Supplementary Figure 4 | S-7 |
| Supplementary Figure 5 | S-7 |
| Supplementary Methods | S-8 |
| -Chemistry | S-8 |
| -Experimental procedures for preparation of pheromones | S-10 |
| -GC analysis with chiral column | S-13 |
| -Copies of NMR spectra | S-16 |
| -References | S-21 |
| List of additional supplementary files | S-21 |

**Supplementary Table 2.** Compounds used for characterization of *Ips typographus* odorant receptors (ORs) and Orco, including their purities, source information, and examples of main biological origins.

| Compound | Purity (%)* | Source* | Examples of biological origin |
|---|---|---|---|
| Acetophenone | 99 | Acros | Beetle, fungi |
| Amitinol | 91 | R. U. | Beetle |
| Anisole | >99 | Sigma-Aldrich | Fungi |
| Benzaldehyde | >99 | Kebo | Non-host, fungi |
| Benzyl acetate | >99 | Aldrich | Fungi |
| Benzyl alcohol | 99 | Aldrich | Non-host, fungi |
| (±)-*exo*-Brevicomin | 99 | W. F. | Beetle, fungi |
| (±)-*endo*-Brevicomin | 96 | Synergy Semiochemicals | Beetle, fungi |
| (±)-Camphene | 95 | Aldrich | Host |
| (±)-Camphor | 97 | Aldrich | Host, fungi |
| (+)-3-Carene | 99 | Aldrich | Host |
| (±)-Carvone | >99 | Fluka | Fungi |
| (±)-Chalcogran | 90 | Celamerck, GmbH | Beetle |
| (±)-1,8-Cineole | 99 | Aldrich | Host |
| Citral (*E*/*Z* mix) | 99 | Aldrich | Fungi |
| (5*S*,7*S*)-*trans*-Conophthorin | 94 | W. F. | Non-host, fungi |
| *p*-Cymene | >99 | Acros | Host |
| 2,3-Dihydrobenzofuran | 99 | Aldrich | Fungi |
| 3,4-Dimethoxytoluene | 98 | Givaudan-Roure | Host, fungi |
| Estragole | >99 | Aldrich | Host, fungi |
| 4-Ethylguaiacol | 98 | Sigma-Aldrich | Fungi |
| Eugenol methyl ether | >99 | Fluka | Host, fungi |
| (±)-Frontalin | >99 | Synergy Semiochemicals | Beetle |
| Geranyl acetate | 97 | Aldrich | Fungi |
| Geranylacetone | >99 | Fluka | Non-host, fungi |
| Hexanal | 96 | Sigma | Non-host |
| 1-hexanol | >99 | Fluka | Non-host, fungi |
| *E*2-hexenal | 98 | Aldrich | Non-host |
| *E*2-hexenol | 96 | Aldrich | Non-host |
| *E*3-hexenol | 98 | Aldrich | Non-host |
| *Z*2-hexenol | 95 | Aldrich | Non-host |
| *Z*3-hexenol | 98 | Aldrich | Non-host |
| (±)-Ipsdienol | 94 | Bedoukian | Beetle |
| (*R*)-(-)-Ipsdienol | 99 (98% ee) | A. M. | Beetle |
| (*S*)-(+)-Ipsdienol | 98 (98% ee) | A. M. | Beetle |
| (±)-Ipsenol | 95 | Synergy Semiochemicals | Beetle |
| (*R*)-(+)-Ipsenol | >99 (99% ee) | A. M. | Not produced by beetles |
| (*S*)-(-)-Ipsenol | >99 (98% ee) | A. M. | Beetle |
| (+)-Isopinocamphone | >99 | R. U. | Host, fungi |
| (-)-Isopinocamphone | >99 | R. U. | Host, fungi |
| Lanierone | >99 | Synergy Semiochemicals | Beetle |
| (-)-Limonene | >99 | Fluka | Host |
| 4-Methyl anisole | >99 | Fluka | Fungi |
| 2-Methyl-1-butanol | >99 | Aldrich | Fungi |
| 3-Methyl-1-butanol | 99 | Aldrich | Fungi |
| 2-Methyl-3-buten-2-ol | >99 | Acros | Beetle, fungi |
| (±)-2-Methylbutyl acetate | >99 | SAFC | Fungi |
| 3-Methylbutyl acetate | 97 | Sigma-Aldrich | Fungi |
| Myrcene | 95 | Sigma-Aldrich | Host |
| *E*-Myrcenol | >99 | Fytofarm | Beetle |
| (±)-Myrtenol | 96 | G. B. | Beetle, fungi |
| (±)-3-Octanol | 97 | Sigma-Aldrich | Non-host, fungi |
| (±)-1-Octen-3-ol | 98 | Janssen Chimica | Non-host, fungi |

| | | | |
|---|---|---|---|
| 2-Phenethyl acetate | >99 | Aldrich | Fungi |
| 2-Phenylethanol | >99 | Sigma | Beetle, fungi |
| (+)-α-Pinene | 98 | Janssen Chimica | Host |
| (-)-α-Pinene | >99 | Fluka | Host |
| (-)-β-Pinene | 92 | Fluka | Host |
| (+)-Pinocamphone | 84 | R. U | Host, fungi |
| (-)-Pinocamphone | 81 | R. U. | Host, fungi |
| (-)-*E*-Pinocarvone | 99 | D. K. | Beetle, fungi |
| (±)-Sabinene | 97 | Chemos GmbH | Host, non-host |
| Styrene | >99 | Fluka | Fungi |
| γ-Terpinene | 97 | Aldrich | Host |
| Terpinolene | 98 | Fluka | Host |
| (±)-4-Thujanol | 97 | Sigma-Aldrich | Host, fungi |
| Toluene | >99 | Merck | Fungi |
| (-)-*cis*-Verbenol | 99 | Borregaard | Beetle |
| (+)-*trans*-Verbenol | 92 | SCM | Beetle |
| (-)-*trans*-Verbenol | 97 | SciTech Ltd., Prague | Beetle |
| (-)-Verbenone | >99 | Fluka | Beetle, fungi |
| 4-Vinyl anisole | 97 | Aldrich | Fungi |
| VUAA1 | 98 | Sigma-Aldrich | None |

* Abbrevations: ee = enantiomeric excess; R. U. = gift from Rikard Unelius (Linnaeus University, Kalmar, Sweden); W. F. = gift from Wittko Francke (University of Hamburg, Germany); A. M. = synthesized in this study by Aleš Machara; G. B. = gift from Gunnar Bergström (University of Gothenburg, Sweden); D. K. = gift from Dineshkumar Kandasamy (Max Planck Institute for Chemical Ecology, Jena, Germany).

**Supplementary Table 3.** Assessment of the completeness of antennal transcriptome Trinity and CLC assemblies using the Benchmarking Universal Single-Copy Orthologs (BUSCOv3) tool performed against the Insecta odb9 dataset (https://busco.ezlab.org/).

| | Trinity | CLC |
|---|---|---|
| Complete BUSCOs (C) | 1615 (97.4%) | 1328 (80.1%) |
| Complete and single-copy BUSCOs (S) | 378 | 1284 |
| Complete and duplicated BUSCOs (D) | 1237 | 44 |
| Fragmented BUSCOs (F) | 27 | 230 |
| Missing BUSCOs (M) | 16 | 100 |
| Total BUSCO groups searched | 1658 | 1658 |

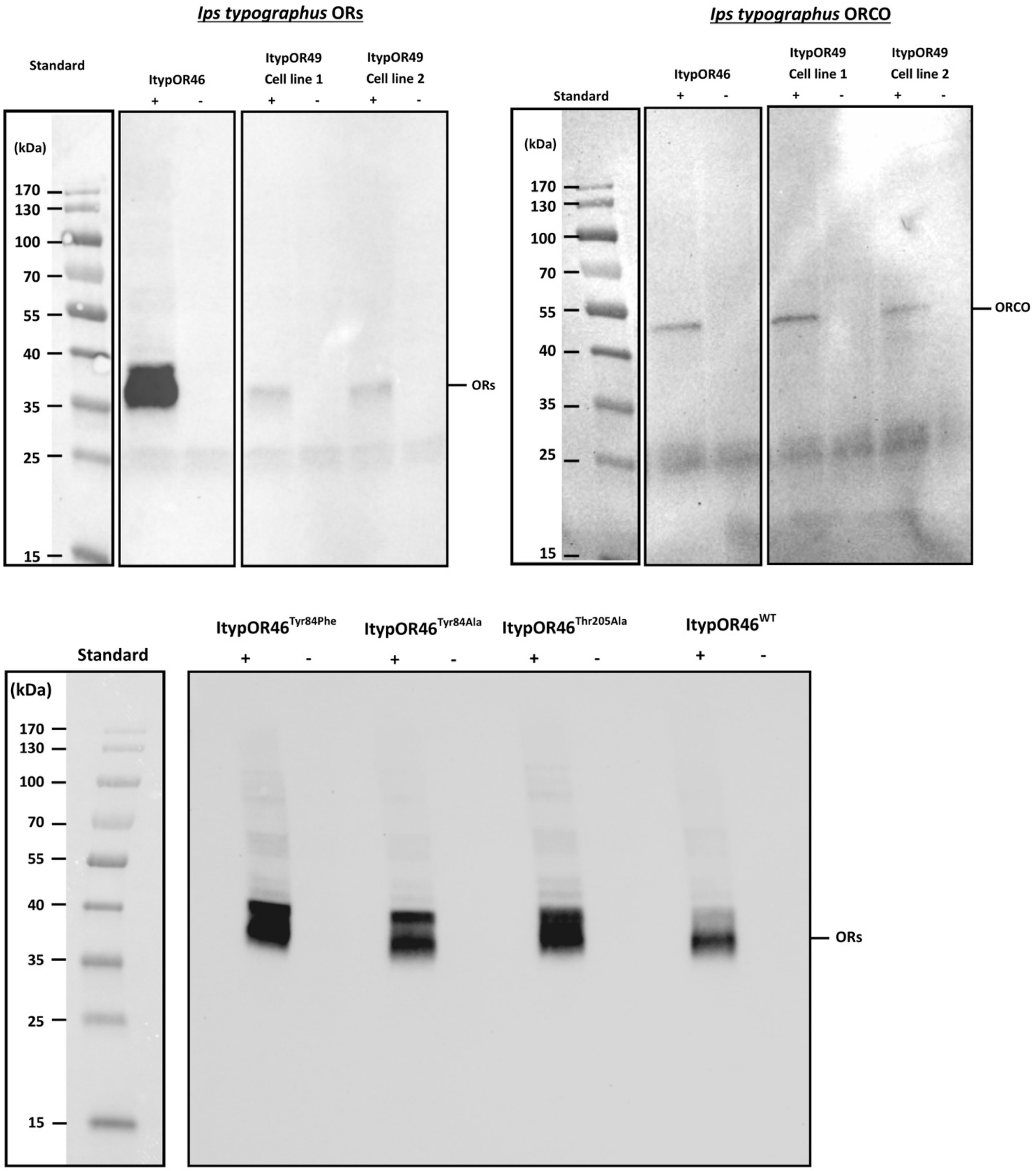


**Supplementary Figure 1.** Protein detection of *Ips typographus* odorant receptors (ORs; V5-tagged) and Orco (myc-tagged) from TREx/HEK293 cells by Western blot. *Upper left panel:* Detection of wildtype ItypOR46 and ItypOR49 (two cell lines). *Upper right panel:* detection of Orco in the same cell lines. *Lower panel:* detection of three versions of mutated ItypOR46 proteins and wildtype (WT) ItypOR46 (included as control). Proteins were only detected from cells induced (+) to express ItypORs and Orco, and not from non-induced (–) control cells, indicating proper regulation by the repression system.

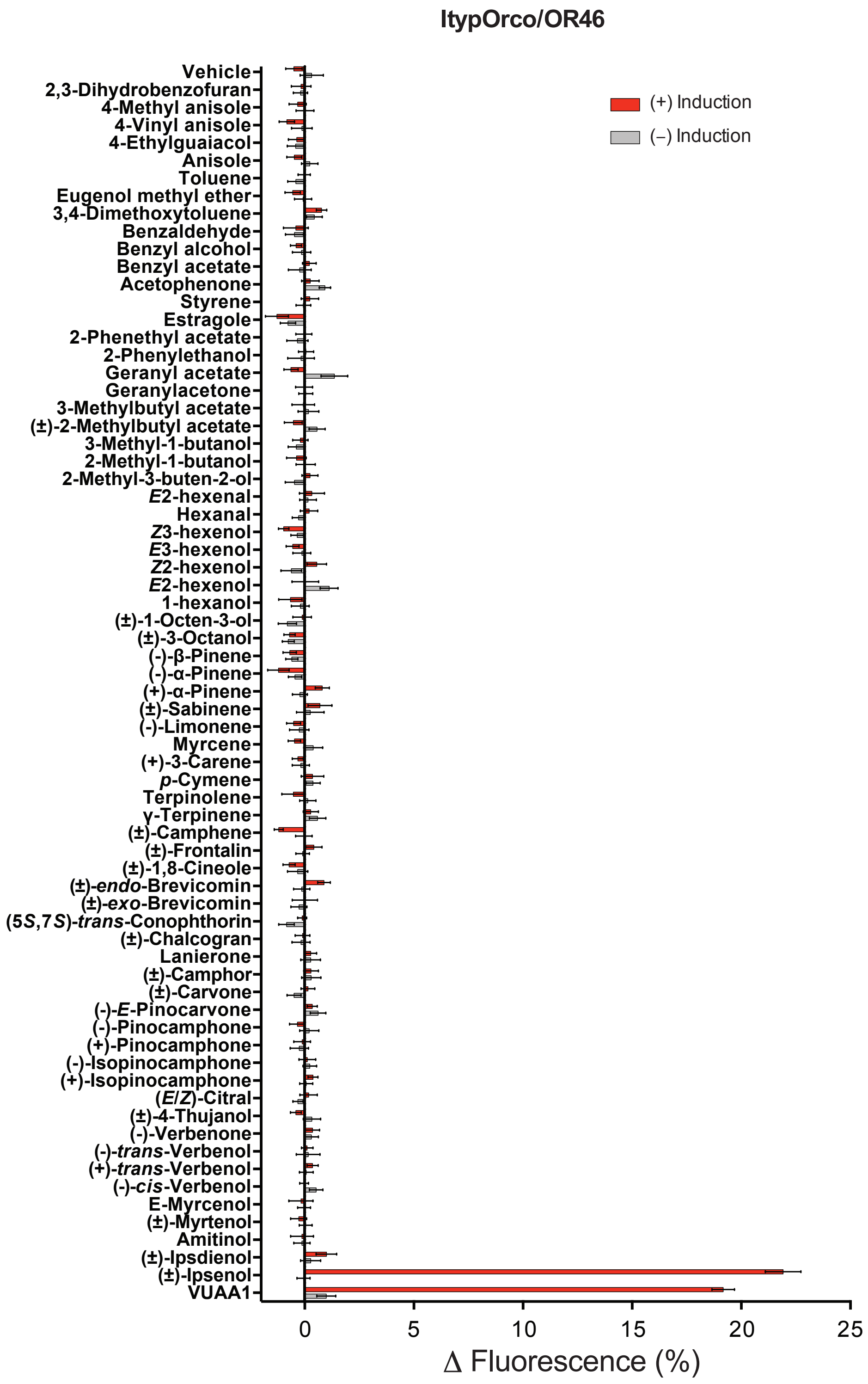


**Supplementary Figure 2.** Response of TREx/HEK293 cells expressing ItypOR46 and ItypOrco to all stimuli (30 μM) and vehicle control in the screening experiment (n = 3 biological replicates, each including 3 technical replicates, i.e., $n_{total} = 9$). (+)-Induction: response of cells induced to express ItypOrco and ItypOR46; (–)-Induction: response of non-induced control cells. Data represent mean responses ± SEM.

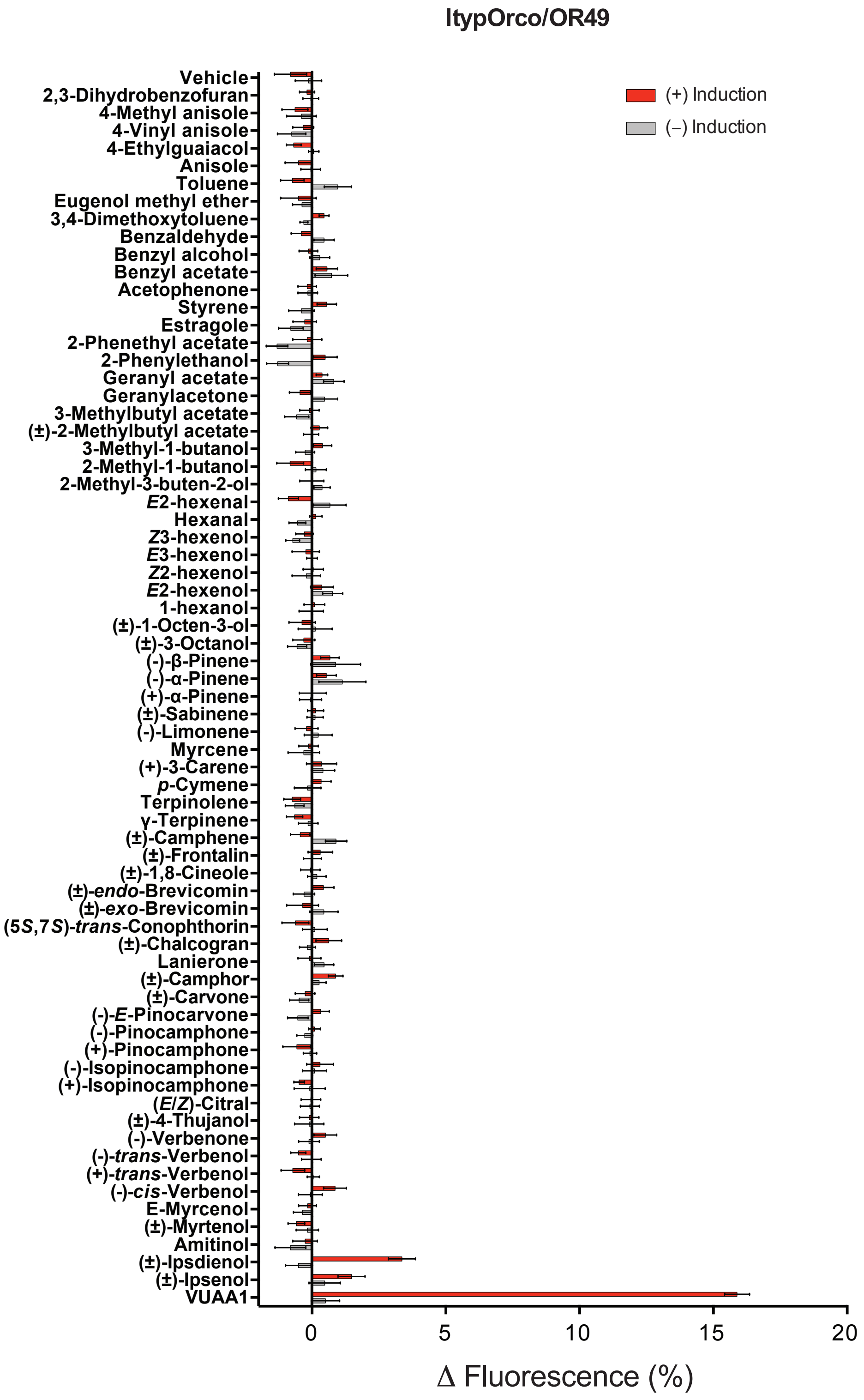


**Supplementary Figure 3.** Response of TREx/HEK293 cells expressing ItypOR49 and ItypOrco to all stimuli (30 μM) and vehicle control in the screening experiment (n = 3 biological replicates, each including 3 technical replicates, i.e., $n_{total}$ = 9). (+)-Induction: response of cells induced to express ItypOrco and ItypOR49; (–)-Induction: response of non-induced control cells. Data represent mean responses ± SEM.

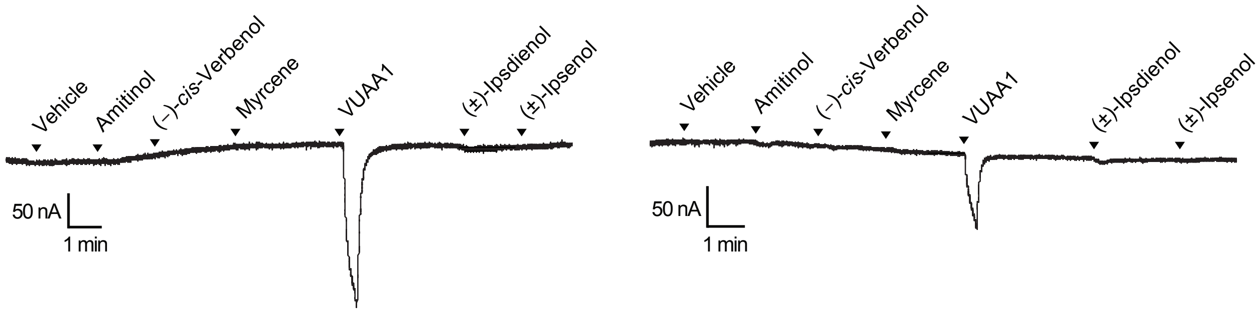


**Supplementary Figure 4.** Current traces of two oocytes expressing ItypOrco/ItypOR49, indicating responses to the Orco agonist VUAA1 and minute responses to racemic ipsdienol.

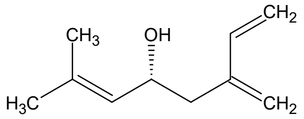

(*R*)-Ipsdienol
(4*R*)-2-methyl-6-methylideneocta-2,7-dien-4-ol

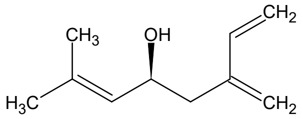

(*S*)-Ipsdienol
(4*S*)-2-methyl-6-methylideneocta-2,7-dien-4-ol

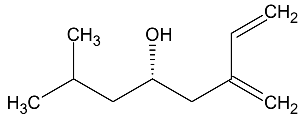

(*S*)-Ipsenol
(4*S*)-2-methyl-6-methylideneoct-7-en-4-ol

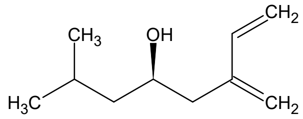

(*R*)-Ipsenol
(4*R*)-2-methyl-6-methylideneoct-7-en-4-ol

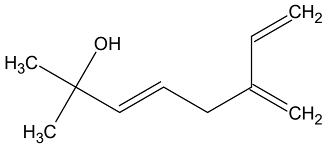

Amitinol
(3*E*)-2-methyl-6-methylideneocta-3,7-dien-2-ol

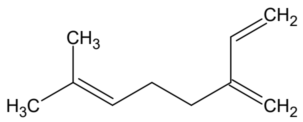

Myrcene
7-methyl-3-methylideneocta-1,6-diene

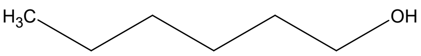

Hexan-1-ol

**Supplementary Figure 5.** Chemical structures and IUPAC names of compounds included in the molecular docking analyses against ItypOR46 and ItypOR49.

### Supplementary Methods

#### Chemistry

Both enantiomers of ipsdienol were obtained by resolution of the corresponding racemate **1** (Scheme 1). The reaction of 2-methylbutenal with isoprenylpotassium[1,2] provided the mentioned racemate **1** that in turn was treated with chiral oxabicyclo[3.3.0]octene[3,4] to get a mixture of separable diastereomers **2a**,**b**. Purified diastereomers **2a** and **2b** were subjected to methanolysis under slightly acidic condition that furnished (*S*)-ipsdienol **1a** and (*R*)-ipsdienol **1b**.

Scheme 1.

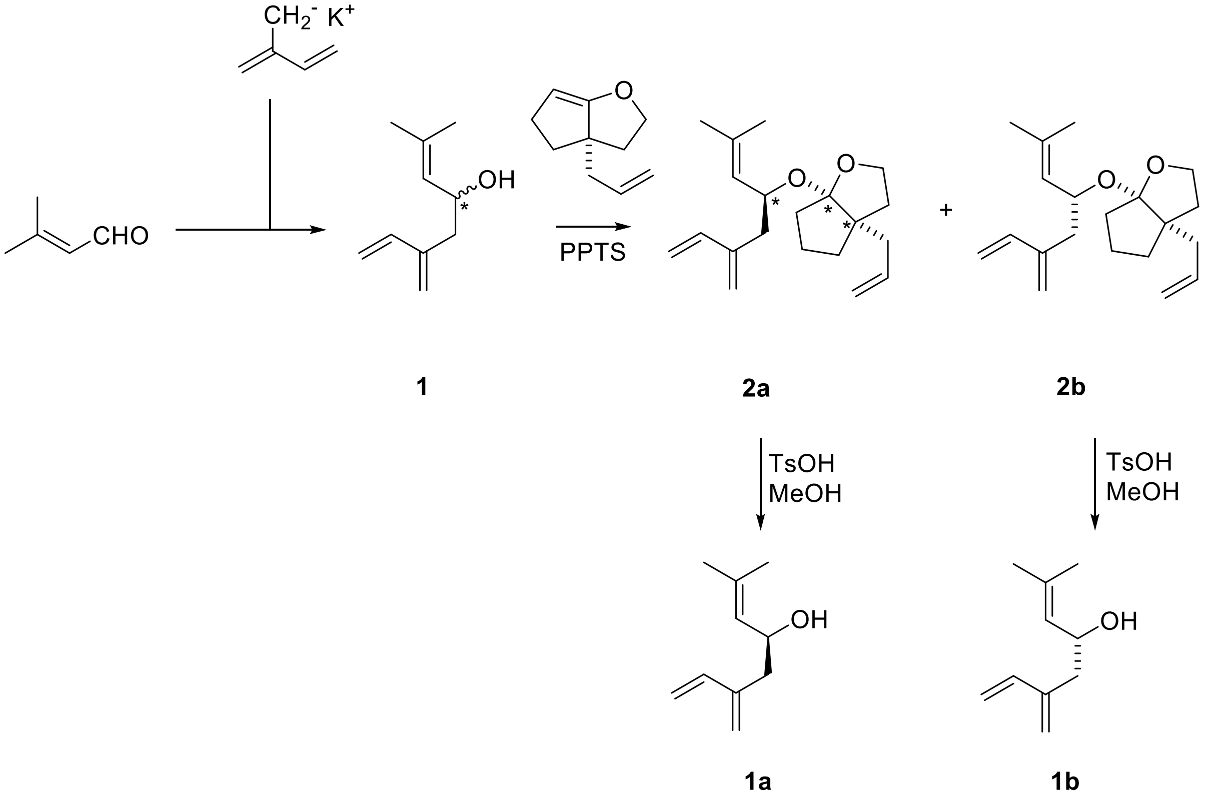


Both (*S*)- and (*R*)-ipsenol (**3a** and **3b**) were prepared with high enantioselectivity (>97% e.e.) following a slightly modified Brown's procedure (Scheme 2).[2] First, metallated isoprene was treated with commercially available chiral *B*-methoxy-diisopinocampheylborane ($Ipc_2BOMe$) to form the "ate" complex, followed by liberation of isoprenylboration reagent using $BF_3.Et_2O$. *In situ*-formed *B*-isoprenyl-diisopinocampheylborane ($Ipc_2BIpn$) was combined with isovaleraldehyde to provide a reaction mixture that was worked-up in a non-oxidative manner:

i.e. simple addition of neat acetaldehyde. The reaction mixture was then concentrated and subsequent column chromatography of the residue afforded ipsenols in good yields.

Scheme 2.

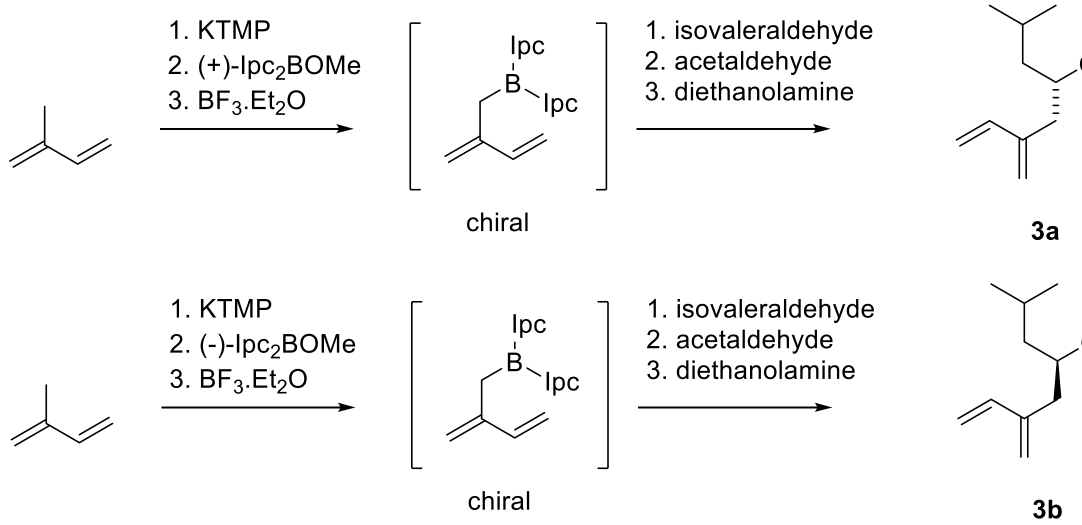


Preparation of racemic ipsenol **3** based on simple addition of isoprenylpotassium to isovaleraldehyde failed, and thus the racemate **3** was prepared in a stepwise fashion (Scheme 3). Isoprenylpotassium was allowed to react with isopropoxy-tetramethyldioxaborolane. The formed isoprenyl boronate **4** was isolated and purified.[5] Its reaction with isovaleraldehyde in toluene afforded racemic ipsenol **3**.

Scheme 3.

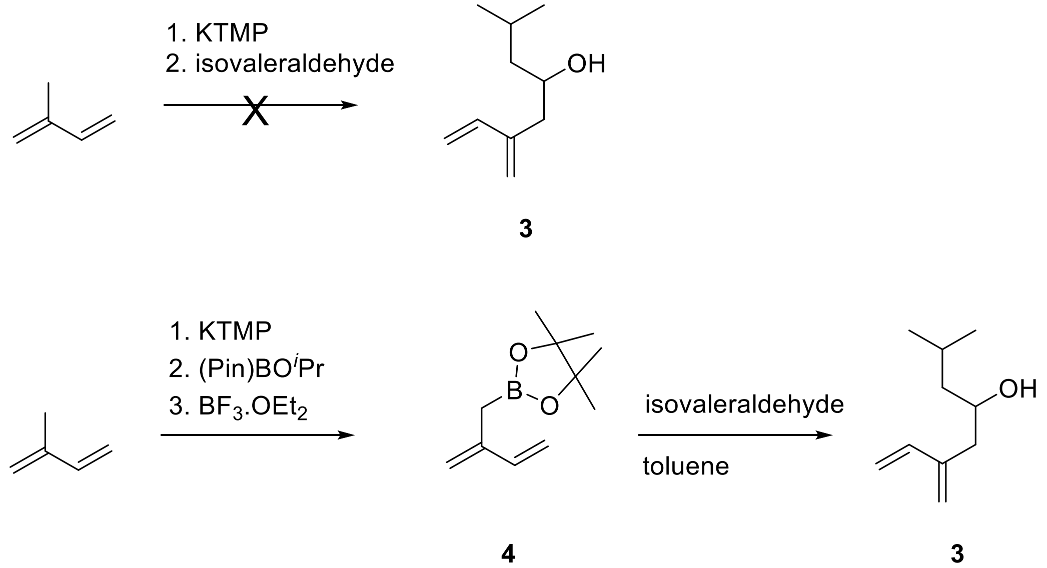

**Experimental procedures**

**General:** Unless otherwise noted, all reactions were carried out under argon using oven-dried glassware. The solvents used for reactions were distilled from drying agents indicated and were transferred under argon: THF (Na/benzophenone); toluene (Na). Chromatography was performed using Fluka silica gel 60 (0.040 - 0.063mm). For TLC analysis, $F_{254}$ –coated aluminum sheets were used. The spots were detected both in UV and by the solution of $Ce(SO_4)_2{\cdot}4H_2O$ (1%) and $H_3P(Mo_3O_{10})_4$ (2%) in 10% sulfuric acid. All starting materials were used as purchased (Sigma Aldrich, TCI), unless otherwise noted. The $^1$H-NMR spectra were measured at 400 MHz, the $^{13}$C-NMR spectra at 100 MHz in $CDCl_3$ with tetramethylsilane or solvent peaks as an internal standard. The chemical shifts are given in *δ*-scale, coupling constants *J* are given in Hz. The EI mass spectra were determined at an ionizing voltage of 70 eV, the *m/z* values are given alone with their relative intensities (%). The ESI mass spectra were recorded using a ZQ micromass mass spectrometer (Waters) equipped with an ESCi multimode ion source and it was controlled by MassLynx software. Methanol was used as solvent.

Racemic ipsdienol **1**

The solution of butyllithium (5 mL, 2.5 M solution in hexane; 12.5 mmol) was added dropwise to a solution of tetramethylpiperidine (1.80 g; 12.5 mmol) in dry THF (5 mL) placed in an ice bath. After 15 min of stirring, the mixture was cooled to -78 °C. A solution of potassium *t*-butoxide (1.40 g; 12.5 mmol) in dry THF (8 mL) was added dropwise. Then solution of 3-methylbutenal (1.05 g; 12.5 mmol) was added slowly and the reaction mixture was removed from the cooling bath and was allowed to reach room temperature. After 30 min the reaction was quenched with a saturated solution of ammonium chloride. Crude product was extracted with diethyl ether. Combined organic layers were washed with water, brine, and concentrated under reduced pressure (120 Torr). Column chromatography (silica gel, eluent DCM) afforded 1.02 g (52%) of colorless liquid. $^1$H NMR (400 MHz, $CDCl_3$) δ 6.60 – 6.23 (m, 1H), 5.45 – 4.96 (m, 5H), 4.54 (td, *J* = 8.4, 4.9 Hz, 1H), 2.63 – 2.27 (m, 2H), 1.76 (s, 3H), 1.73 (s, 3H).

Resolution of racemic ipsdienol, preparation of compounds **2a** and **2b**

A mixture of racemic ipsdienol (0.62 g; 4.07 mmol), (*S*)-5-allyl-2-oxabicyclo[3.3.0]oct-8-ene (0.73 g; 4.88 mmol) and catalytic amount of pyridinium *p*-toluenesulfonate (approx. 0.002 g) in anhydrous DCM (5 mL) was stirred for 5 hours at room temperature. Afterwards the mixture was concentrated under reduced pressure. The obtained residue was subjected to thorough column chromatography (silica gel, eluent toluene: cyclohexane/2:3) that yielded two diastereomers. Diastereomer **2a**- 0.4 g (32%), $[\alpha]_D$ = -11.1° (c = 0.38, MeOH). $^1$H NMR (400 MHz, $CDCl_3$) δ 6.38 (dd, *J* = 17.6, 10.8 Hz, 1H), 5.88 (ddt, *J* = 17.3, 10.1, 7.3 Hz, 1H), 5.36 (d, *J* = 17.6 Hz, 1H), 5.21 – 4.89 (m, 6H), 4.66 (dt, *J* = 9.1, 6.8 Hz, 1H), 3.92 – 3.58 (m, 2H), 2.51 (dd, *J* = 13.4, 7.1 Hz, 1H), 2.38 – 2.19 (m, 2H), 2.17 – 2.03 (m, 1H), 1.99 – 1.85 (m, 2H), 1.70 (m, 4H), 1.62 – 1.50 (m, 8H). $^{13}$C NMR (100 MHz, $CDCl_3$) δ 143.2, 138.9, 137.2, 131.9, 128.5, 118.2, 117.2, 116.5, 113.4, 68.8, 65.7, 54.5, 40.3, 39.7, 38.4, 36.2, 35.3, 25.7, 21.7, 18.4. HR-CI-MS *m/z* calculated for $C_{20}H_{30}O_2$ $[M+H]^+$ 303.2324, found: 303.2328.

Diastereomer **2b**- 0.4 g (32%), $[\alpha]_D$ = -94.2° (c = 0.26, MeOH). $^1$H NMR (400 MHz, $CDCl_3$) δ 6.38 (dd, *J* = 17.6, 10.7 Hz, 1H), 5.87 (ddt, *J* = 17.3, 10.1, 7.3 Hz, 1H), 5.28 (d, *J* = 17.6 Hz, 1H), 5.20 – 4.84 (m, 6H), 4.57 – 4.38 (m, 1H), 3.90 – 3.48 (m, 2H), 2.46 (dd, *J* = 13.7, 7.7 Hz, 1H), 2.36 – 2.20 (m, 2H), 2.15 – 1.99 (m, 1H), 1.92 (ddd, *J* = 11.8, 7.2, 4.3 Hz, 1H), 1.78 – 1.32 (m, 12H). $^{13}$C NMR (100 MHz, $CDCl_3$) δ 143.1, 139.1, 137.2, 130.6, 128.7, 118.7, 117.6, 116.4, 113.2, 69.9, 65.5, 54.6, 40.4, 39.3, 38.2, 36.6, 34.7, 25.7, 21.4, 18.3.

Preparation of (*S*)-ipsdienol **1a**

A solution of compound **2a** (0.1 g; 0.33 mmol) and catalytic amount of pyridinium *p*-toluenesulfonate (approx. 0.005 g) in anhydrous methanol (2 mL) was stirred for five days. The reaction mixture was diluted with diethyl ether (5 mL) and washed with water. The organic layer was concentrated under reduced pressure and the residue was purified by column chromatography (silica gel, eluent DCM). Methanolysis afforded 0.03 g (58%) of the product. Chiral GC determined the enantiomeric excess to be 98.2%. $[\alpha]_D$ = +13.1° (c = 0.18, MeOH).

Preparation of (*R*)-ipsdienol **1b**

This compound was prepared using a procedure similar to the one used for the preparation of the enantiomer **1a**. Methanolysis of **2b** (0.1 g; 0.33 mmol) afforded 0.025 g (49%) of the title compound. Chiral GC determined the enantiomeric excess to be 98.0%. $[\alpha]_D$ = -13.2° (c = 0.19, MeOH).

Preparation of (*S*)-ipsenol **3a**

The solution of butyllithium (5 mL, 2.5 M solution in hexane; 12.5 mmol) was added dropwise to a solution of tetramethylpiperidine (1.80 g; 12.5 mmol) in dry THF (5 mL) placed in an ice bath. After 15 min of stirring the mixture was cooled to -78 °C. A solution of potassium *t*-butoxide (1.40 g; 12.5 mmol) in dry THF (8 mL) was added dropwise. Then neat isoprene (1.30 g; 18.0 mmol) was added slowly to the mixture, which turned red. The reaction mixture was stirred at -60 °C for 20 min. Afterwards the mixture was cooled to -78 °C and a solution of (+)-*B*-methoxy-diisopinocampheylborane (3.95 g; 12.5 mmol) in THF (3 mL) was added. The reaction mixture was allowed to stir for 20 min at -78 °C and then neat boron trifluoride etherate (2.0 mL; 16.0 mmol) was added dropwise in 5 min. Isovaleraldehyde (1.1 g; 12.5 mmol) in THF (3 mL) was added dropwise to a rapidly stirred mixture of $Ipc_2BIpn$, maintained at -78 °C. Neat acetaldehyde (1.1 mL) was added in one portion and the mixture was allowed to warm to 0 °C. THF was evaporated on the rotavap and the obtained residue was diluted with diethyl ether (20 mL). Finally, diethanolamine (1.3 mL; 14.0 mmol) was added. The mixture was stirred for 1 hour. The formed suspension was passed through a plug of Celite. The filtrate was concentrated under reduced pressure and chromatographed (silica gel; gradient elution started with straight pentane, then was used mixture of pentane:DCM/ 9:1 to 1:1). The *one pot* reaction afforded 0.7 g (36%) of colorless liquid. $[\alpha]_D$ = -13.0° (c = 0.29, MeOH). Chiral GC technique determined enantiomeric excess as 97.8%. HR-MS-TOF-EI+ *m/z* calculated for $C_{10}H_{18}O$ $[M]^+$ 154.1358, found: 154.1362.

Preparation of (*R*)-ipsenol **3b**

This compound was prepared using a procedure similar to the one used for the preparation of the enantiomer **3a**. The *one pot* reaction using of (-)-*B*-methoxy-diisopinocampheylborane (3.95 g; 12.5 mmol) afforded 0.8 g (41%) of colorless liquid. $[\alpha]_D$ = +10.4° (c = 0.27, MeOH). Chiral GC determined the enantiomeric excess to be 98.8%.

Preparation of isoprenylpinacolboronic ester **4**

The solution of butyllithium (10 mL, 2.5 M solution in hexane; 25.0 mmol) was added dropwise to a solution of tetramethylpiperidine (3.54 g; 25.0 mmol) in dry THF (10 mL) placed in an ice bath. After 15 min of stirring the mixture was cooled to -78 °C. A solution of potassium *t*-

butoxide (2.80 g; 25.0 mmol) in dry THF (15 mL) was added dropwise. Then neat isoprene (2.60 g; 37.0 mmol) was added slowly to the mixture, which turned red. The reaction mixture was stirred at -60 °C for 20 min. Afterwards the mixture was cooled to -78 °C and 2-isopropoxy-4,4,5,5-tetramethyldioxaborolane (4.6 g; 25.0 mmol) was added. After 20 min the reaction was quenched by addition of boron trifluoride etherate (4.0 mL; 32.0 mmol). The reaction mixture was concentrated down to thick oil that was loaded onto silica gel. Column chromatography (eluent cyclohexane:$Et_2O$/ 10:1) afforded 1.86 g (38%) of colorless oil. $^1$H NMR (400 MHz, $CDCl_3$) δ 6.47 (ddd, *J* = 17.4, 10.5, 0.6 Hz, 1H), 5.20 (dd, *J* = 17.4, 0.8 Hz, 1H), 5.11 – 4.99 (m, 3H), 1.89 (s, 2H), 1.26 (s, 12H). $^{13}$C NMR (100 MHz, $CDCl_3$) δ 143.1, 139.8, 116.6, 113.7, 83.4, 24.7.

Racemic ipsenol **3**

A mixture of boronate **4** (0.65 g; 3.35 mmol) and isovaleraldehyde (0.29 g; 3.35 mmol) in dry toluene (4 mL) was stirred for two days at room temperature. Then the reaction was quenched with a saturated solution of ammonium chloride (2 mL). The product was extracted with diethyl ether. Combined organic layers were washed with brine and gently concentrated (the ether was evaporated). Column chromatography (eluent cyclohexane: $Et_2O$/ 2:1) afforded 0.31 g (60%) of the desired product as colorless liquid. $^1$H NMR (400 MHz, $CDCl_3$) δ 6.42 (dd, *J* = 17.6, 10.7 Hz, 1H), 5.31 – 4.96 (m, 4H), 3.85 (tt, *J* = 8.7, 4.0 Hz, 1H), 2.52 (ddd, *J* = 13.9, 3.6, 1.1 Hz, 1H), 2.23 (dd, *J* = 13.9, 9.1 Hz, 1H), 1.85 (dddd, *J* = 13.2, 12.1, 8.6, 6.6 Hz, 1H), 1.48 (ddd, *J* = 13.7, 8.5, 5.3 Hz, 1H), 1.30 (ddd, *J* = 13.6, 8.6, 4.2 Hz, 1H), 0.97 (d, *J* = 6.7 Hz, 3H), 0.95 (d, *J* = 6.6 Hz, 3H). $^{13}$C NMR (100 MHz, $CDCl_3$) δ 143.2, 138.5, 118.5, 114.2, 67.6, 53.4, 46.5, 40.6, 24.7, 23.5, 22.1.

**Gas chromatographic (GC) analysis with chiral column**

All analyses were carried out using gas chromatography coupled with mass spectrometric detection (GC-MS) using a TRACE 1310 GC with an ISQ LT mass spectrometer with electron ionization (70 eV), equipped with a quadrupole mass analyzer (Thermo Scientific, Waltham, MA, USA) and a Cyclodex-B column (30 m, i.d. 0.25 mm, 0.25 μm film thickness, Agilent, Santa Clara, CA, USA). The temperature program was 90 °C (80 min) to 200 °C (10 min) at 15

°C/min. The inlet temperature was set to 200 °C and the samples were introduced in split mode (1:30). Helium was used as a carrier gas, set to a constant flow at 1 ml/min.

(*S*)-ipsdienol **1a**

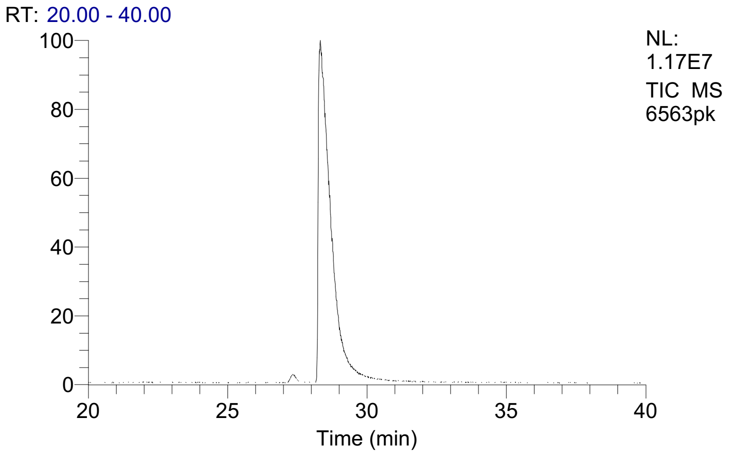


| % Area |
|---|
| 0.9 |
| 99.1 |

(*R*)-ipsdienol **1b**

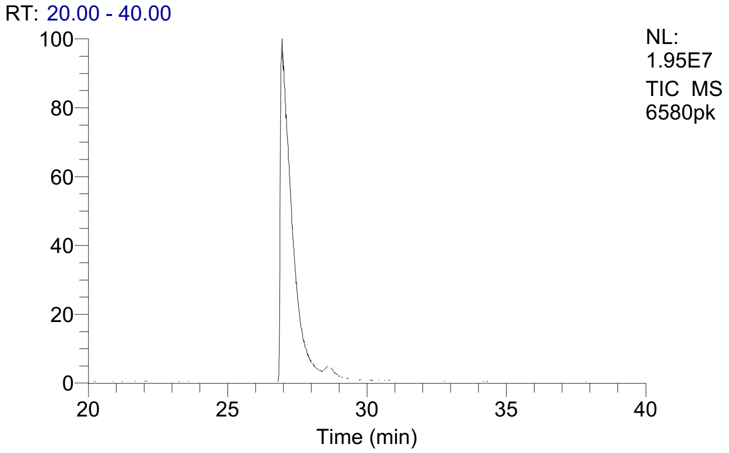


| % Area |
|---|
| 99.0 |
| 1.0 |

(*S*)-ipsenol **3a**

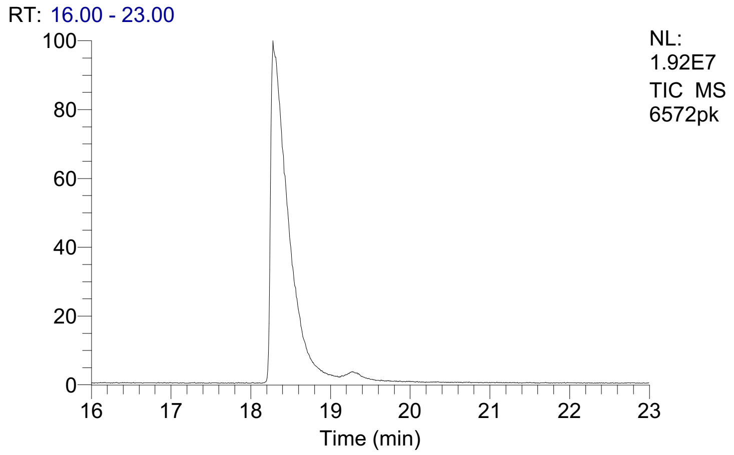


| % Area |
|---|
| 98.9 |
| 1.1 |

(*R*)-ipsenol **3b**

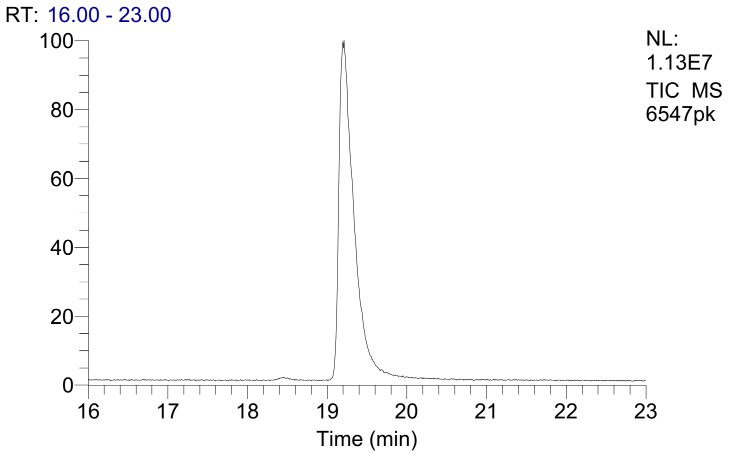


| % Area |
|---|
| 0.6 |
| 99.4 |

**Copies of NMR spectra**

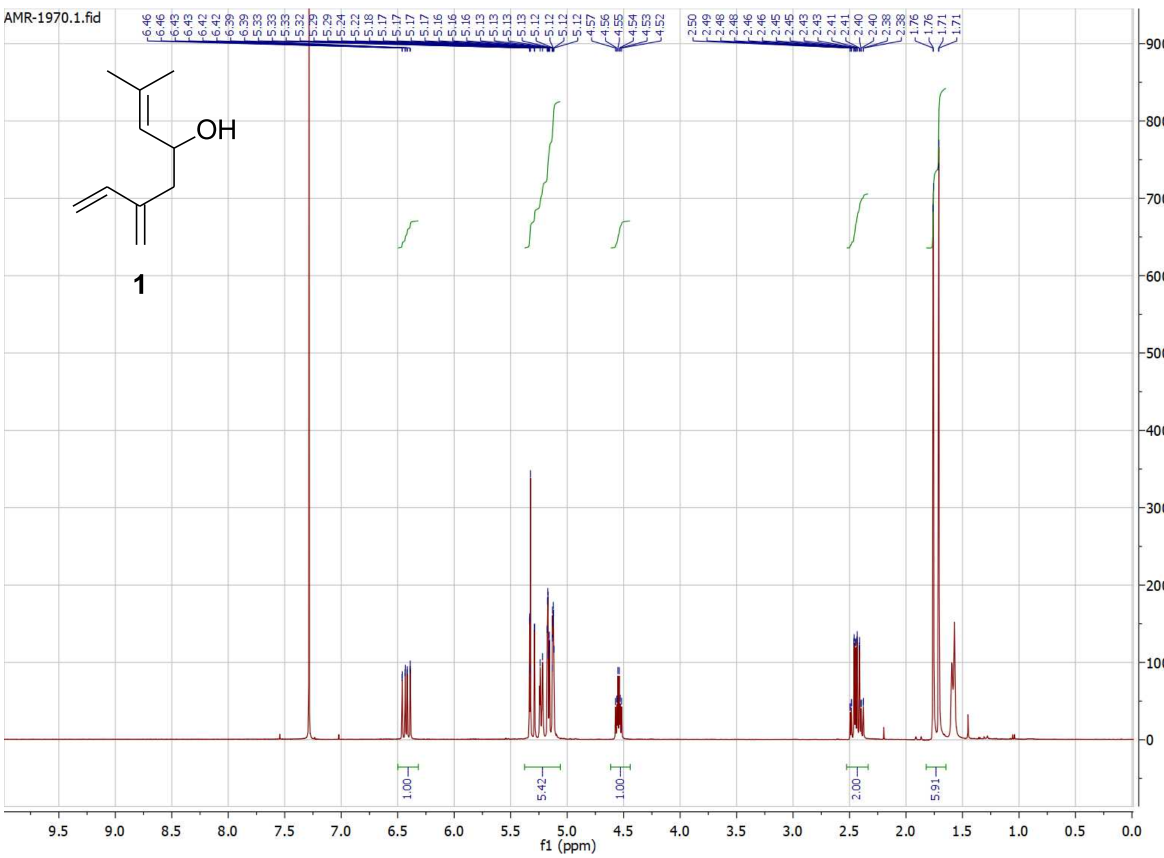

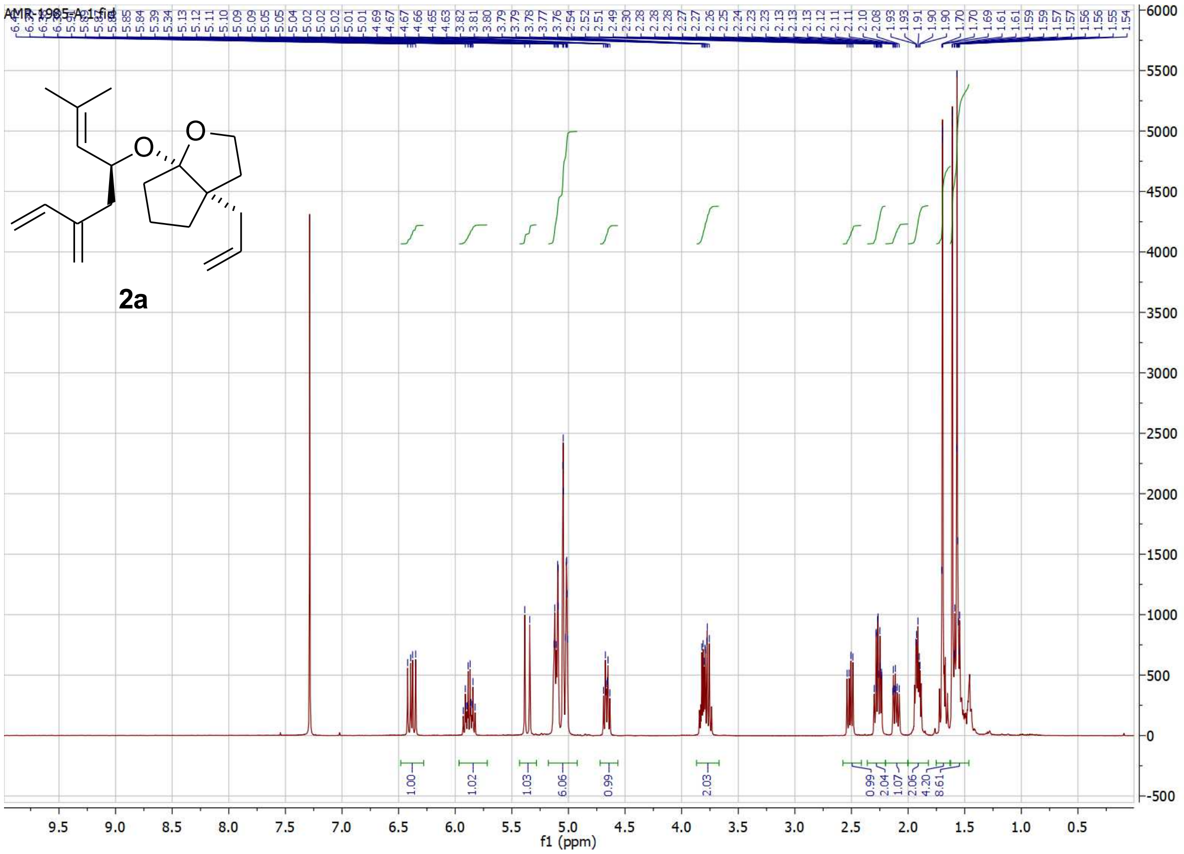
AMR-1985-A.1.fid
2a
1.00
1.02
1.03
6.06
0.99
2.03
0.99
2.04
1.07
2.06
4.20
8.61
f1 (ppm)

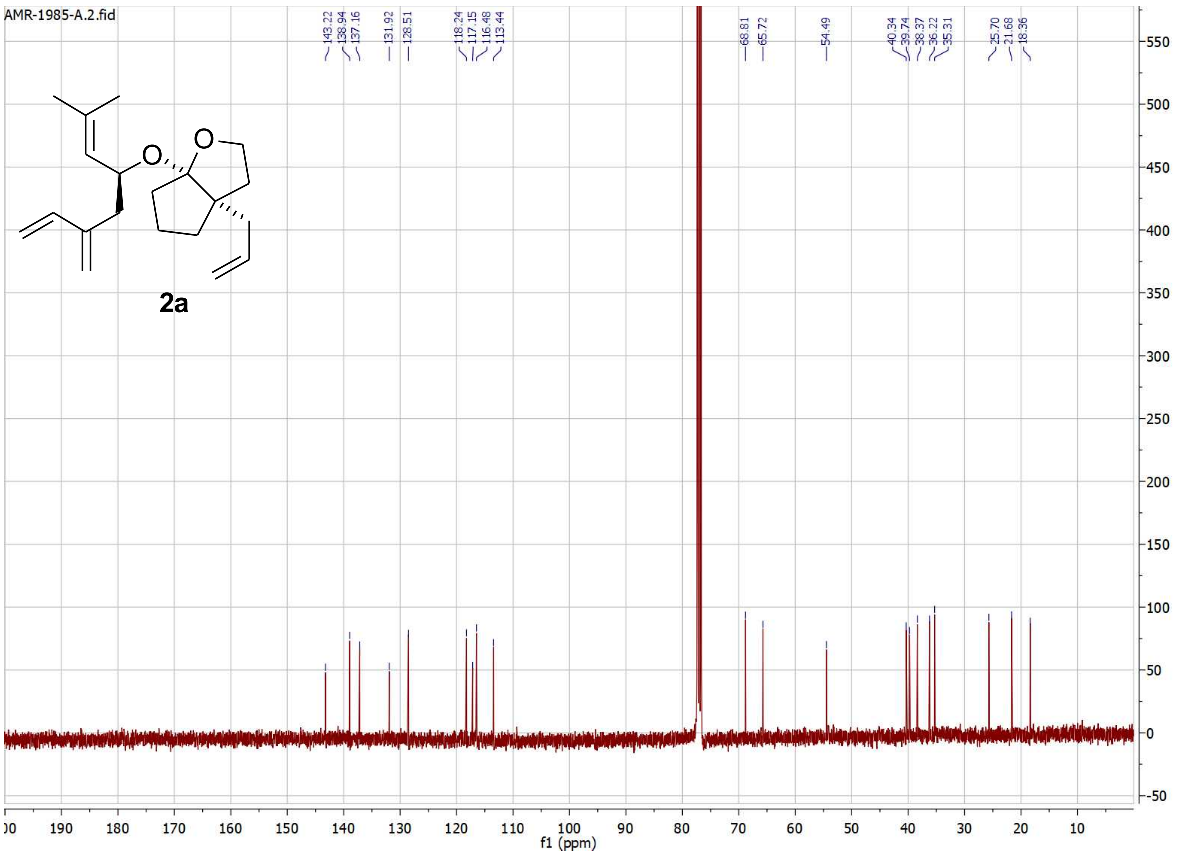
AMR-1985-A.2.fid
143.22
138.94
137.16
131.92
128.51
118.24
117.15
116.48
113.44
68.81
65.72
54.49
40.34
39.74
38.37
36.22
35.31
25.70
21.68
18.36
2a
f1 (ppm)

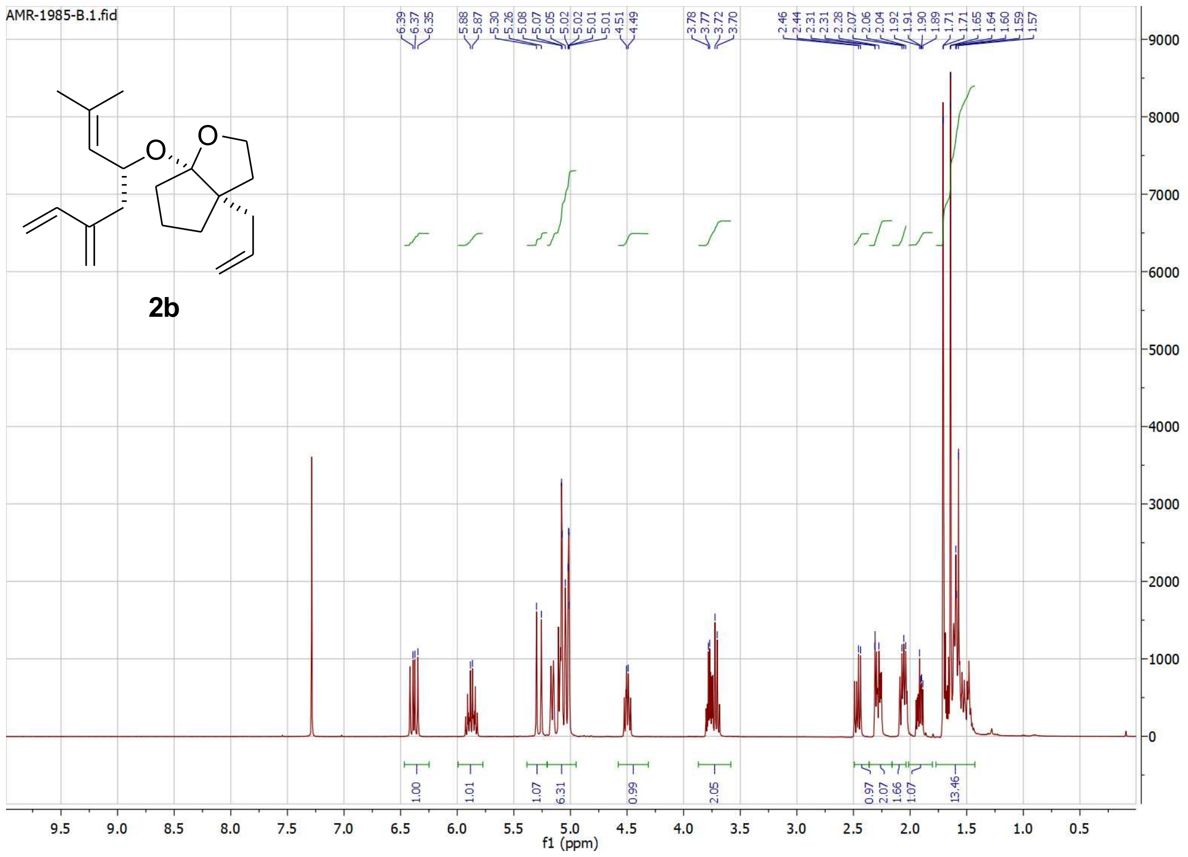
AMR-1985-B.1.fid
2b
f1 (ppm)

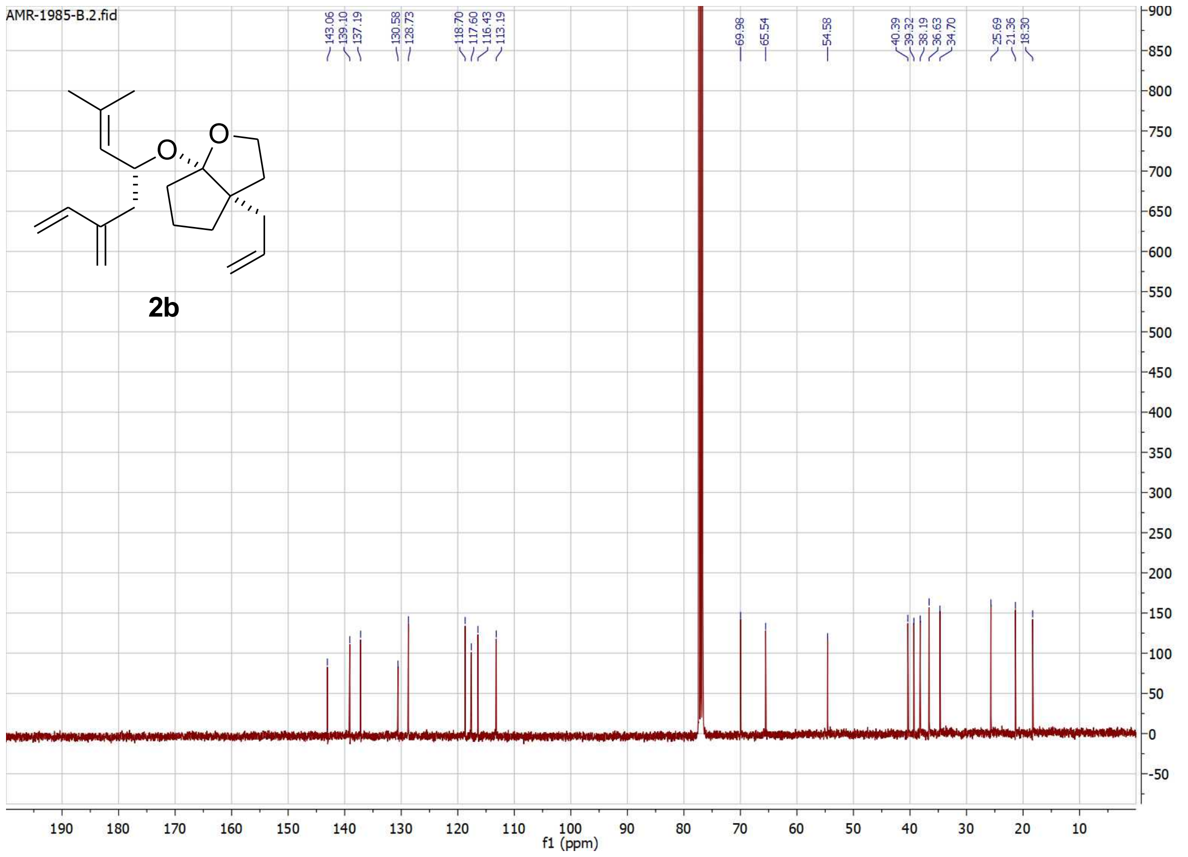
AMR-1985-B.2.fid
2b
f1 (ppm)

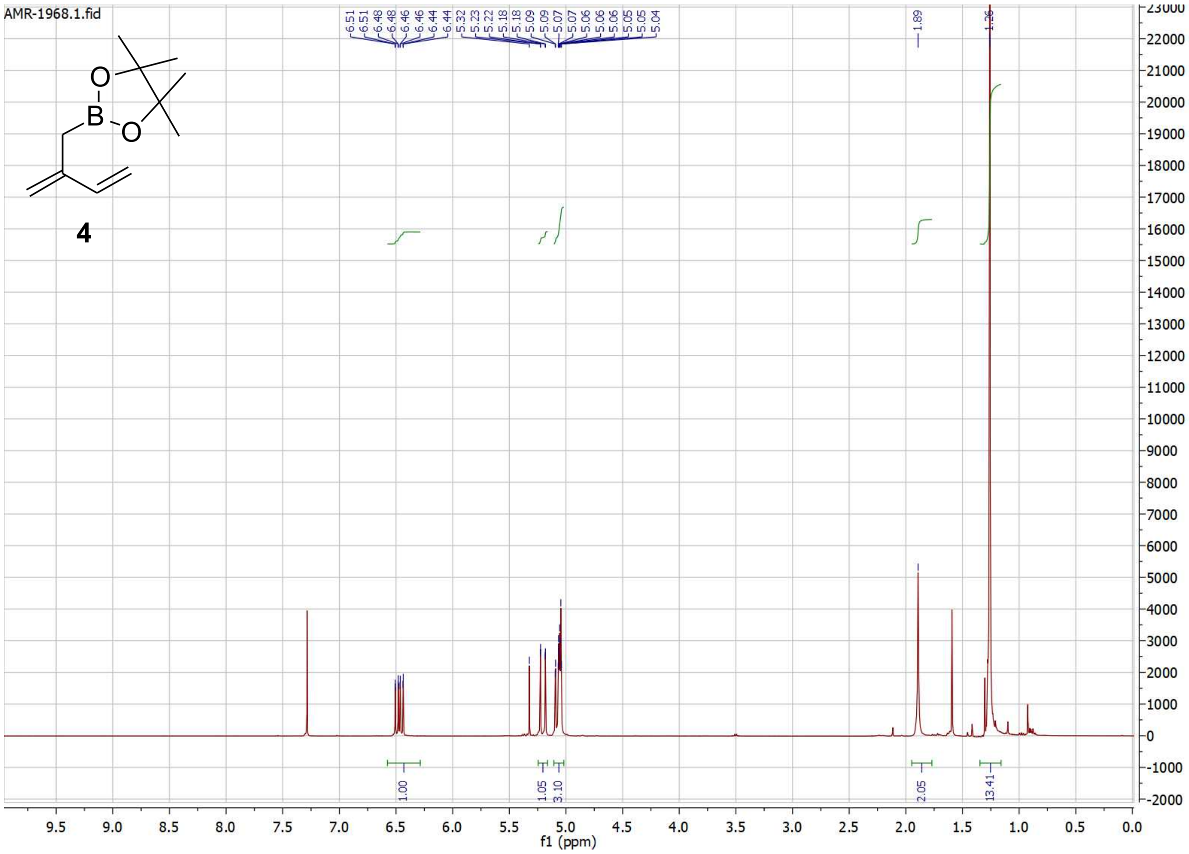
AMR-1968.1.fid
4
f1 (ppm)

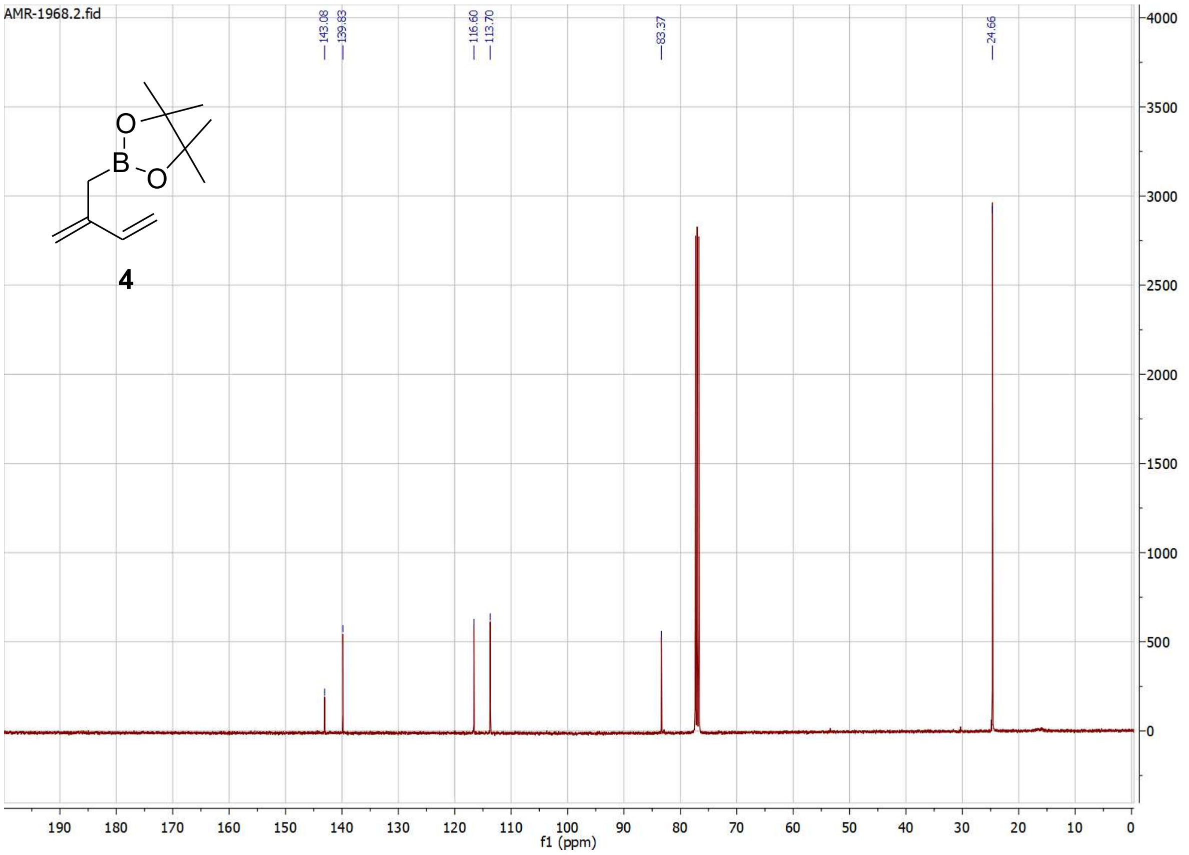
AMR-1968.2.fid
4
f1 (ppm)

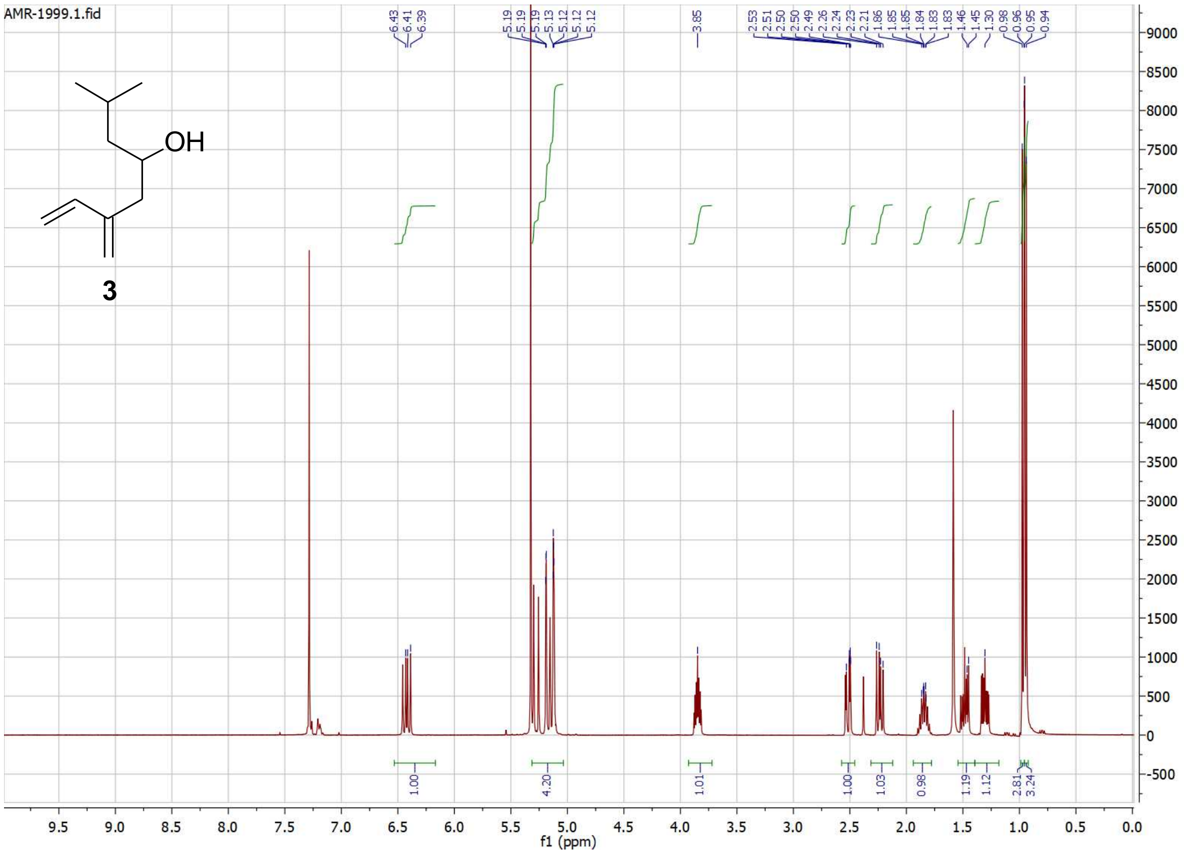
AMR-1999.1.fid
OH
3
6.43
6.41
6.39
5.19
5.19
5.19
5.13
5.12
5.12
5.12
3.85
2.53
2.51
2.50
2.50
2.49
2.26
2.24
2.23
2.21
1.86
1.85
1.85
1.84
1.83
1.83
1.46
1.45
1.30
0.98
0.96
0.95
0.94
1.00
4.20
1.01
1.00
1.03
0.98
1.19
1.12
2.81
3.24
f1 (ppm)

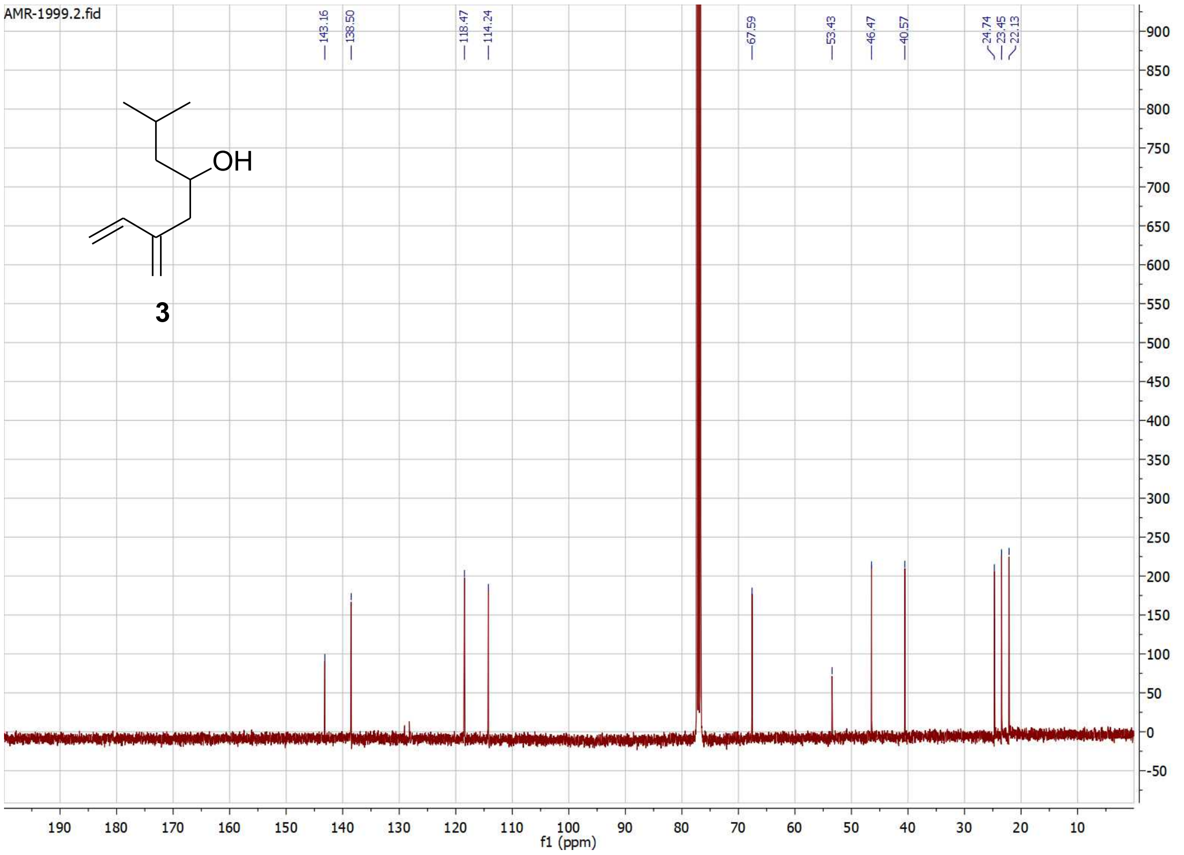
AMR-1999.2.fid
OH
3
143.16
138.50
118.47
114.24
67.59
53.43
46.47
40.57
24.74
23.45
22.13
f1 (ppm)

### List of additional supplementary files

Supplementary Table 1

Supplementary Data 1
